# Brain network dynamics reflect psychiatric illness status and transdiagnostic symptom profiles across health and disease

**DOI:** 10.1101/2025.05.23.655864

**Authors:** Carrisa V. Cocuzza, Sidhant Chopra, Ashlea Segal, Loïc Labache, Rowena Chin, Kaley Joss, Avram J. Holmes

## Abstract

The network organization of the human brain dynamically reconfigures in response to changing environmental demands, an adaptive process that may be disrupted in a symptom-relevant manner across psychiatric illnesses. Here, in a transdiagnostic sample of participants with (n=134) and without (n=85) psychiatric diagnoses, functional connectomes from intrinsic (resting-state) and task-evoked fMRI were decomposed to identify constraints on brain network dynamics across six cognitive states. Hierarchical clustering of 110 clinical, behavioral, and cognitive measures identified participant-specific symptom profiles, revealing four core dimensions of functioning: internalizing, externalizing, cognitive, and social/reward. Brain network dynamics were flattened across cognitive states in individuals with psychiatric illness and could be used to accurately separate dimensional symptom profiles more robustly than both case-control status and primary diagnostic grouping. A key role of inhibitory cognitive control and frontoparietal network interactions was uncovered through systematic model comparison. We provide novel evidence that brain network dynamics can accurately differentiate the extent that psychiatrically-relevant dimensions of functioning are exhibited across health and disease.

## Introduction

The human brain is a complex dynamical system^1,2^ that enables rapid, context-relevant information processing to occur in a time-varying manner^3,4^. This fundamental property of brain functioning enables flexible and adaptive responses within brain regions and across large-scale functional networks^5–15^. Substantial progress has been made in characterizing topological properties that allow large-scale brain networks to maintain a balance between functional flexibility and the stability required for central nervous system integrity^7,16,17^. For example, intrinsic (i.e., resting-state)^18^ functional connectivity has been characterized in terms of its small worldness^19,20^, hub structure^21,22^, hierarchical organization^23–25^, multiple network assignments^26–28^, and probabilistic distributions^29^. This literature provides converging evidence for the key role of network-based mechanisms in the facilitation of flexible information processing across the cortical sheet^7,16,30^. While resting-state connectivity patterns reliably approximate the properties of functional specialization within and between large-scale brain systems^31–33^, task-linked changes in connectivity patterns are thought to enable dynamic shifts in context-dependent processing over time^16,34,35^. Moreover, there is mounting evidence that time-varying, flexible shifts in connectivity patterns (here termed *reconfigurations*^30,36–39^) more optimally account for individual differences in both cognition and behavior than traditional, “static” approaches for studying brain functions^7,40–44^. Importantly, given that flexible processing is key for adaptively responding to changing environmental or cognitive demands, alterations in brain network dynamics are theorized to contribute to the onset and maintenance of psychiatric illness^45–49^. However, despite the importance of network reconfigurations in accounting for individual differences in behavior across health and disease, the extent that brain network dynamics link with psychiatric symptom structure remains unclear.

Disrupted brain dynamics may constitute a key mechanism that accounts for symptom profiles across patient populations. This is evident, for example, in patients diagnosed with schizophrenia, where electrophysiological studies have shown disrupted oscillatory activity and transient synchronizations, particularly in low-frequency bands^4,50,51^. Recent work has uncovered dynamic connectivity patterns in patients diagnosed with bipolar disorder and schizophrenia that reflect symptoms of mania and psychosis^52–56^. Suboptimal network dynamics have also been reported in obsessive compulsive disorder^57^, attentional disorders^58^, and major depressive disorder^59,60^. Importantly, with respect to predicting diagnostic status, network dynamics (**Fig. 1A**, e.g., inter- or across-state changes in connectivity, as in **Fig. 1B**) tend to outperform measures that assume stable, temporally-invariant, patterns of connectivity^53,58,61^. However, despite growing evidence for altered network dynamics in patient populations, the extent that shifts in connectivity patterns across cognitive states accounts for individual- specific symptom structure remains to be established. Here, we examine brain-based features that constrain across-state network reconfigurations, and how such dynamics may be impaired in a symptom-relevant manner across psychiatric illnesses (**Fig. 1B**).

**Figure 1.**
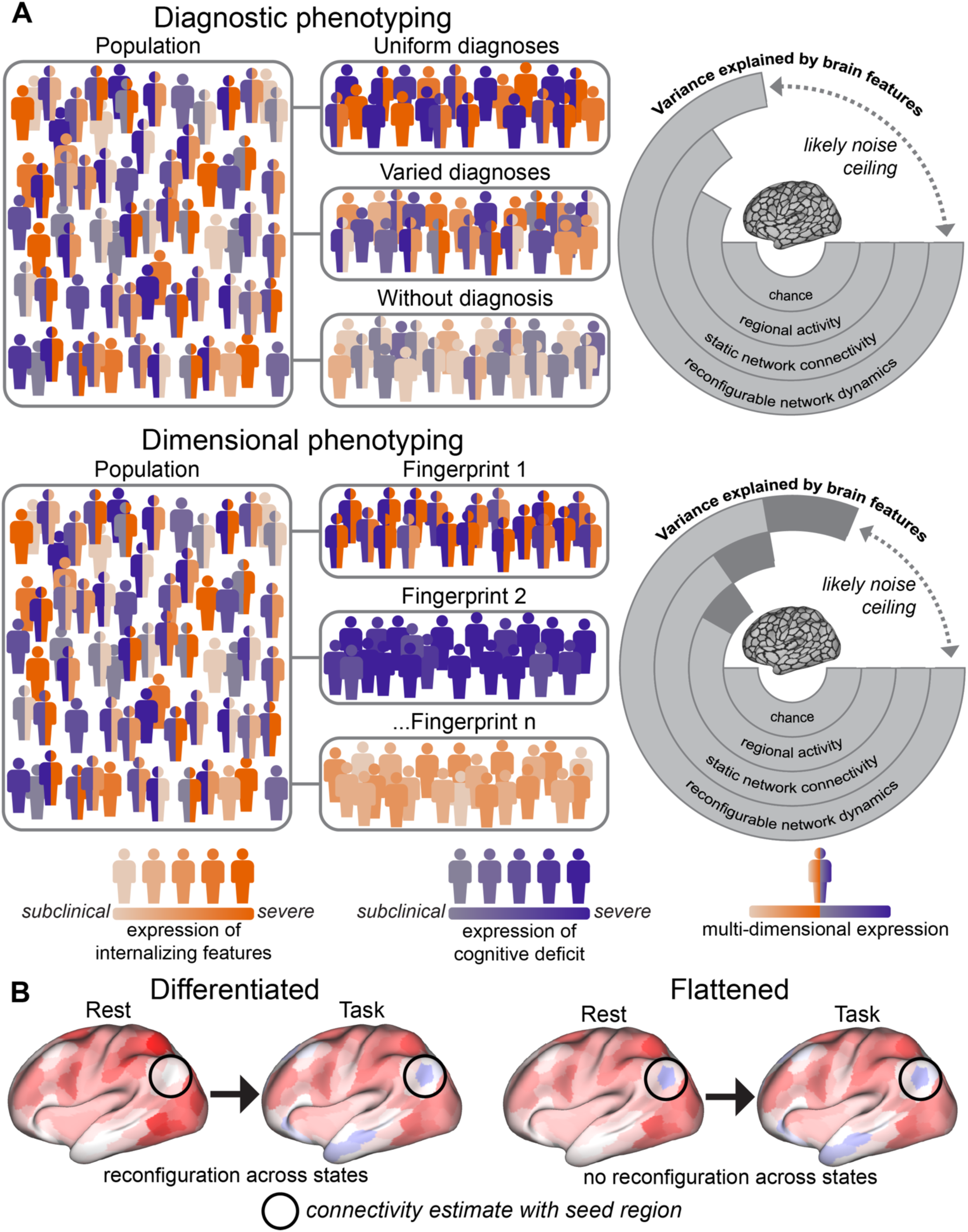
Theoretical framework. (**A**) Schematic distinguishing diagnostic (top) and dimensional phenotypes (bottom), with respect to establishing brain-behavior relationships. Color saturations: symptom expression levels in two example dimensions of functioning: internalizing and cognition. Uniform diagnoses: example individuals with the same diagnosis but variable symptom expression. Varied diagnoses: different diagnoses but overlapping symptoms. Without diagnoses: subclinical or transient expression. For simplicity, individuals with zero symptoms are not shown. Dimensional phenotyping groups individuals based on symptom expression patterns (fingerprints), which are thus highly consistent in each group. We propose that, generally, neurobiological features better account for the dimensionally-based phenotypes (right circle plots), given increased consistency in underlying symptom structure. Further, it is likely that some brain features link with behavior better than others; with brain network dynamics outperforming static network features, univariate brain activity, and chance. Noise ceiling: theoretical cap on explainable variance with current neuroimaging, although this may increase with future methods. (**B**) Example schematic of how clinically-relevant brain network dynamics may be exhibited across cognitive states, for example, shifting from rest to task states. In health, connectivity patterns shift in a small but functionally-relevant manner to optimally meet task demands. Here, there is a differentiable change in connectivity that flexibly and adaptively supports the online recruitment of processing resources. In less healthy systems, one possibility is that rest-to-task shifts in connectivity are suboptimal such that little-to-no reconfiguration (flattened, or less differentiated) leads to failures in recruiting the resources needed to account for task demands or behavioral goals.

One longstanding challenge in establishing reliable links between brain processes and clinical presentation is the observation that psychiatric symptoms tend to co-occur across discrete diagnostic categories, while the expression of symptoms within a given diagnosis group can also be highly variable^62–70^ (**Fig. 1A**). For instance, patients diagnosed with either major depressive disorder or schizophrenia can present with internalizing symptoms, such as anhedonia, avolition, fatigue, and flat affect^71,72^. There may also be overlapping expressions of cognitive deficits across diagnoses, such as a reduced ability to plan, impaired working memory, and a tendency to ruminate^73,74^. Moreover, individuals not meeting criteria for diagnosis can also express subclinical or transient functional deficits. For example, a non-diagnosed individual may temporarily experience low levels of mood, motivation, or reward sensitivity. Observations such as this have prompted some theorists to reframe psychopathology as a hierarchy of deficits across broad dimensions (also termed domains) of cognitive and behavioral functioning, rather than a collection of distinct diagnostic constructs^75–80^, emphasizing the importance of individualized, precision medicine approaches^81,82^.

Here, we adopt a transdiagnostic framework by analyzing data from a sample of individuals with and without psychiatric diagnoses. We used cognitive and behavioral measures to uncover individually- profiled symptom structures across core dimensions of psychiatrically-relevant functioning^62,68,69,83–85^ (**Fig. 1**), which we term symptom fingerprints. Given the importance of brain network dynamics to adaptive information processing, we hypothesized that reconfiguring connectivity patterns across a varied set of cognitive states would accurately separate individuals based on dimensionally-organized symptom structure^3,53,58,61^ (**Fig. 1**), as well as by their case-control status and primary diagnostic labels. We used non-negative matrix factorization (NMF) to model brain network reconfiguration dynamics, which can reveal underlying constraints upon connectivity patterns as individuals transition across cognitive states^86–92^. We discovered that network reconfiguration dynamics were flattened (i.e., less differentiated) across cognitive states in patients, and these dynamics could be used to accurately predict case-control status, primary diagnoses, and participant-specific symptom fingerprints. The link between network dynamics and dimensionally-organized symptom fingerprints was prominently driven by shifts from intrinsic to cognitive control^93,94^ (i.e., Stroop^95,96^) task states. Further, control (frontoparietal), thalamic, and task-linked (i.e., visual and somato/motor) network regions were particularly important for classifying symptom fingerprints.

## Results

### Brain network dynamics are flattened in psychiatric illness

We applied non-negative matrix factorization (NMF) to functional connectivity (FC) estimates from each participant and cognitive state to characterize across-state brain network dynamics (**Fig. 2A-C**). NMF has previously identified transition dynamics between states with varied cognitive demand in healthy adults^86^. NMF decomposes positively-thresholded inputs (here: across-state FC patterns) into two lower rank matrices (**Fig. 2C**). These matrices can be multiplied to approximately reconstruct the input connectivity patterns, and each extract distinct, but related, attributes of input connectivity patterns. One is a features matrix, which is similar to a basis set – here termed subgraphs. When applied to an across- state connectivity input, subgraph values quantify features constraining brain network dynamics^86,97^. The next matrix contains coefficients, analogous to an encoding matrix. Coefficients quantify the expression of each subgraph over each observation, which here were cognitive states and participants. Consistent with other dimensionality reduction techniques, NMF extracts a hidden structure that best accounts for a given input dataset. Compared to other techniques, such as PCA, NMF has the benefit of being parts- based, allowing for straightforward inferences regarding reconstruction accuracy. NMF is also less constrained with respect to orthogonality. Non-orthogonality is neurobiologically relevant for multi-state or multi-modal data, as features underlying time-varying neural data are likely expressed with some overlap. For example, while we explicitly examined reconfigurations across intrinsic and task-evoked FC patterns, these states share a structural connectivity backbone^98–100^ and their underlying dynamics are not strictly orthogonal. NMF-uncovered subgraph features, and corresponding expressions of those subgraphs, allowed us to infer key constraints upon the dynamics that underlie across-state brain network reconfigurations. Cross-validation of the discovery dataset tuned NMF parameters as follows: alpha (regularization)=0.2, beta (loss)=1.2, and k (number of subgraphs)=5 (**Supplemental Fig. S1**).

**Figure 2.**
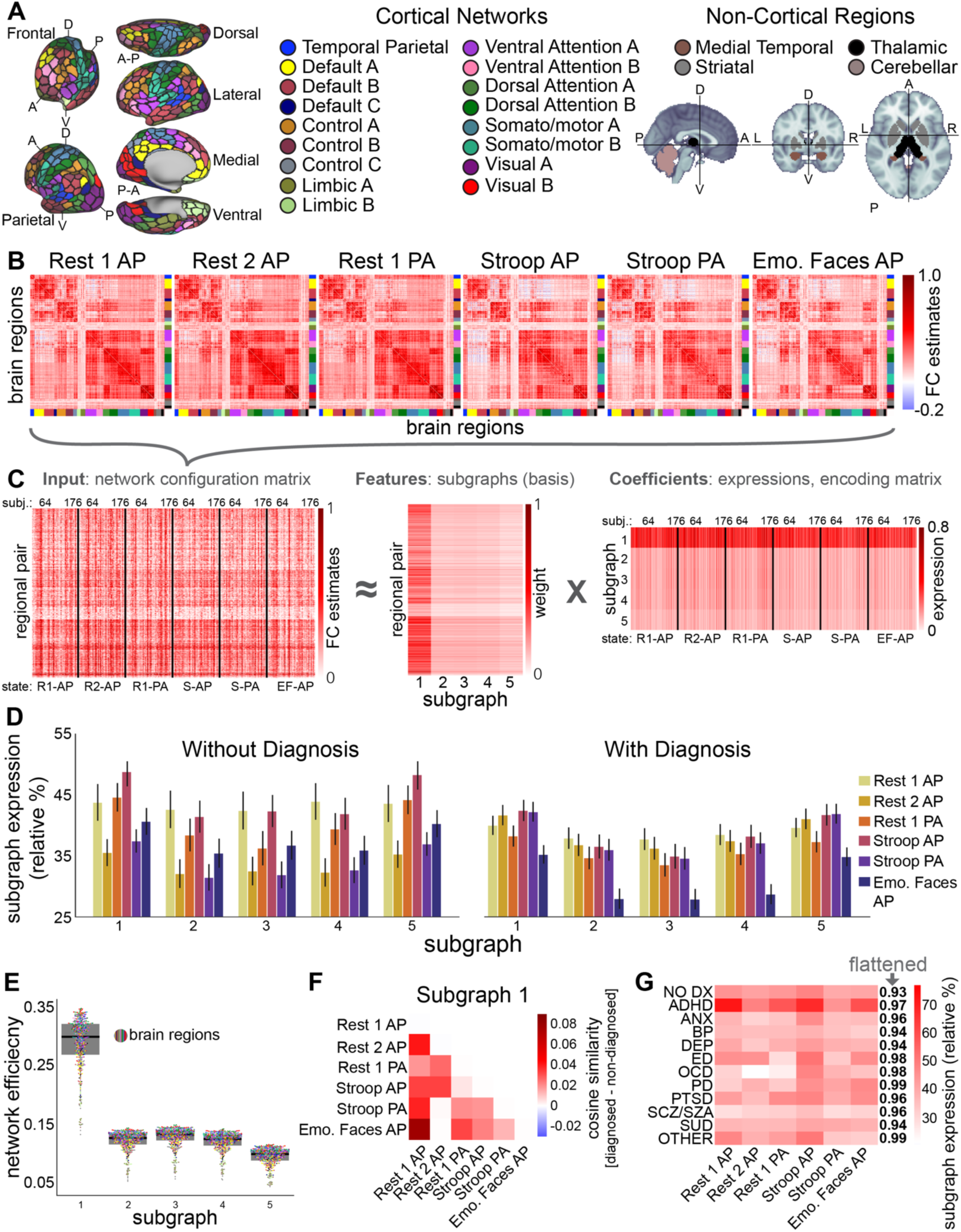
Across-state brain network dynamics in health and psychiatric illness. (**A**) The organization of the human cerebral cortex is revealed through patterns of intrinsic functional connectivity and anatomical segmentation. Left: network organization of the human cortex was based on the 17- network solution from Yeo et al.^105^ across the 400-parcel atlas from Yan et al.^106^ (black borders around cortical regions). Control network = frontoparietal cognitive control network. Right: non-cortical regions were based on Tian et al.^107^ (subcortex) and Buckner et al.^108^ (cerebellum). Network color-coding is used in all subsequent figures. (**B**) FC estimates (across-participant average r, Fisher-z transformed) for each of the six fMRI scans (cognitive states). Regions (x, y axes) sorted per network assignment. Grey arrow: FC was used as input to NMF in panel C. “Region” refers to functional parcellation of the cortex or anatomical subdivisions of non-cortex, and the term “network” refers to a collection of such regions with underlying organization, for simplicity (see **Methods** for details). (**C**) NMF pipeline. Input across-state network configuration matrix (left): FC estimates from upper triangle of B (positively thresholded); flattened and stacked vertically, by state and participant (black lines). NMF decomposes this into subgraph features (middle) and expression coefficients (right). (**D**) Across-participant average expression coefficients (C, right; normalized relative percent) for each subgraph and state. Here, participants were grouped by case-control status. (**E**) Local efficiency of subgraph features (C, middle), projected back into region-by-region networks. Dots: brain regions (averaged across participants; grand average via black line) color-matched for networks. Subgraph 1 exhibited the most efficiency, suggesting information flows across these features more readily. We highlight subgraph 1 based on this (and NMF orthogonality) in all downstream analyses (2-5 subgraph results in Supplement). (**F**) Across-participant cosine similarity of expression coefficients (D) were computed for each group and pair of states. Non-diagnosis-group similarities were subtracted from diagnosis-group similarities. Network dynamics were more flattened (less differentiated) across participants with psychiatric diagnoses in most state-to-state comparisons. (**G**) Same as F, but participants grouped by primary diagnosis (rows). NO DX: no diagnosis; ANX: anxiety; BP: bipolar disorder; DEP: depression; ED: eating disorder; PD: panic disorder or phobia; SCZ/SZA: schizophrenia/schizoaffective; SUD: substance use disorder. Rightmost column: average cosine similarity (here, an index flattened dynamics) of each diagnostic group (i.e., averaging the lower triangle in F, without group-based subtraction).

While we tested hypotheses about dimensionally-organized symptom profiles downstream (see following **Results**), we first tested the hypothesis that network dynamics are relatively flattened in participants with psychiatric diagnoses (**Fig. 1**). The across-state expression patterns of subgraph coefficients (**Fig. 2C**) were less differentiated (i.e., less variable, or more flattened, across states) in those with psychiatric diagnoses relative to those without diagnoses (**Fig. 2D**). Given that across-state subgraph coefficient patterns were expressed consistently in each of the k=5 subgraphs for both groups (**Fig. 2D**), we first restructured each subgraphs’ features (**Fig. 2C**, middle matrix; **Supplemental Fig. S1**) into standard region-by-region networks (**Fig. 2A-B**) and then applied a network efficiency metric^101–103^. Subgraph 1 exhibited the greatest network efficiency (**Fig. 2E**) both locally (brain regions) and globally (functional brain networks), suggesting that information flows across this subgraph’s functional architecture more readily than subgraphs 2 through 5. Thus, for ease of interpretation, main-text **Results** refer to subgraph 1 and subgraphs 2 through 5 are reported in the **Supplement**.

Next, we quantified the extent that network dynamics were flattened for subgraph 1 (subgraphs 2-5: **Supplemental Figs. S1-S2**) by applying cosine similarity to the vectors of participants’ expression coefficients for each state-to-state pair. Thus, a higher cosine similarity score indicates that subgraph features underlying FC reconfiguration dynamics were expressed similarly across cognitive states. We subtracted without-diagnosis similarity scores from with-diagnosis similarity scores for each pair of states (**Fig. 2F**). Thus, a positive difference in cosine similarity indicated relative flattening of dynamics, which we expected for non-diagnosed participants relative to diagnosed participants (**Fig. 1B**). In the majority of state-to-state pairings, the transdiagnostic psychiatric sample exhibited more flattened subgraph expression coefficients. Across state-to-state pairs, the average cosine similarity of those without diagnosis was 0.91 (standard deviation (*SD*)=0.01) and 0.95 with diagnoses (*SD*=0.03; cosine similarity can range from -1 to 1). The difference of with-diagnosis minus without-diagnosis cosine similarities across all state-to-state pairs was significantly greater than zero (*t*(14)=5.08, *p*=8.38 x 10^-5^), suggesting that the dynamics constraining across-state brain network reconfigurations were comparatively flattened in those with psychiatric diagnoses versus those without diagnoses.

Next, we explored the extent that primary diagnostic groups^104^ exhibited flattened brain network dynamics (**Fig. 2G**). The most flattened dynamics were exhibited by those diagnosed with panic disorder, eating disorder, obsessive compulsive disorder, and attention-deficit/hyperactivity disorder (and “other”, which had n=2 therefore we caution against over-interpretation). A moderate amount (in this sample) of flattened brain network dynamics was exhibited by those diagnosed with an anxiety disorder, bipolar disorder, a depressive disorder, schizophrenia/schizoaffective disorder, posttraumatic stress disorder, and substance use disorder. Dynamics were flattened (given by cosine similarity, **Fig. 2G**) to a similar extent in varied diagnostic groups. This is consistent with the literature suggesting that neurobiological estimates may be noisy across select diagnostic categories (**Fig. 1A**) ^67^. To further account for this, we examined the relationship between brain network reconfiguration dynamics and dimensions of functioning underlying psychiatric illness in remaining sections.

*Symptom fingerprints: core dimensions of functioning are exhibited across health and psychiatric illness* Known limitations in identifying robust links between psychiatrically-relevant behavior and the brain include heterogeneous symptomatology within diagnostic categories as well as overlapping symptom structures across diagnostic groups^67,68,84^ (**Fig. 1A**). One meta-analysis reported that resting-state networks exhibited alterations common to more than six distinct diagnostic categories^109^. This suggests that prior methods were either poorly constrained with respect to differentiating diagnostic groups (and likely more applicable to transdiagnostic symptom expression), or that static-network-based biomarkers may lack sensitivity to a single diagnostic group (or both). Here, we accounted for two key constructs for psychiatric phenotyping. First, by using the TCP dataset^110^, which includes participants with and without varied mental health concerns and diagnoses. Second, we incorporated behavioral data with wide coverage across dimensions of functioning (**Supplemental Table S1**). This strategy aims to improve validity by optimizing the balance between precise, individualized phenotyping and broad, group-level diagnostic bins.

After curating, transforming, and imputing 110 behavioral measures that covered a wide breadth of cognitive, behavioral, and psychiatric domains (henceforth referred to collectively as *behavioral measures* for simplicity) (**Supplemental Table S1** and **Figs. S3, S4**), we performed hierarchical clustering^80,111^ on the resulting matrix of individual differences correlations (**Fig. 3A**). The optimal number of clusters according to 18 performance indices was four (**Fig. 3A** black boxes), which is consistent with prior work examining the latent structure of various behaviors^80,112,113^. To limit researcher bias in naming these clusters, we pre-labeled each measure with possible dimensions of functioning (full mapping: **Supplemental Table S1, Fig. S4**). Following a separate PCA on the observed scores of each of the four clusters of measures, we identified which dimensions of functioning was dominant in each cluster by the largest percentage of labels, which were weighted by factor loadings on the first PC. Accordingly, the four clusters were labelled: internalizing (approximately 40% of measures in the cluster had this as the primary label), externalizing (31% of measures), cognition (27%), and social/reward (20%). Canonical examples of measures in the internalizing cluster included (**Fig. 3B**) the Depression, Anxiety, and Stress Scales^114^ and the neuroticism factor of the NEO Five-Factor Inventory-3 (NEO-FFI-3^115^). Example externalizing measures included the extraversion factor of the NEO-FFI-3 and the Young Mania Rating Scale^116^ irritability subscale. Example cognition measures included the entire TestMyBrain suite^117^ and Shipley intelligence measures^118^. Example social/reward measures included the Experiences in Close Relationships^119^ scales and the Temperament and Character Inventory^120^ novelty seeking subscale.

**Figure 3.**
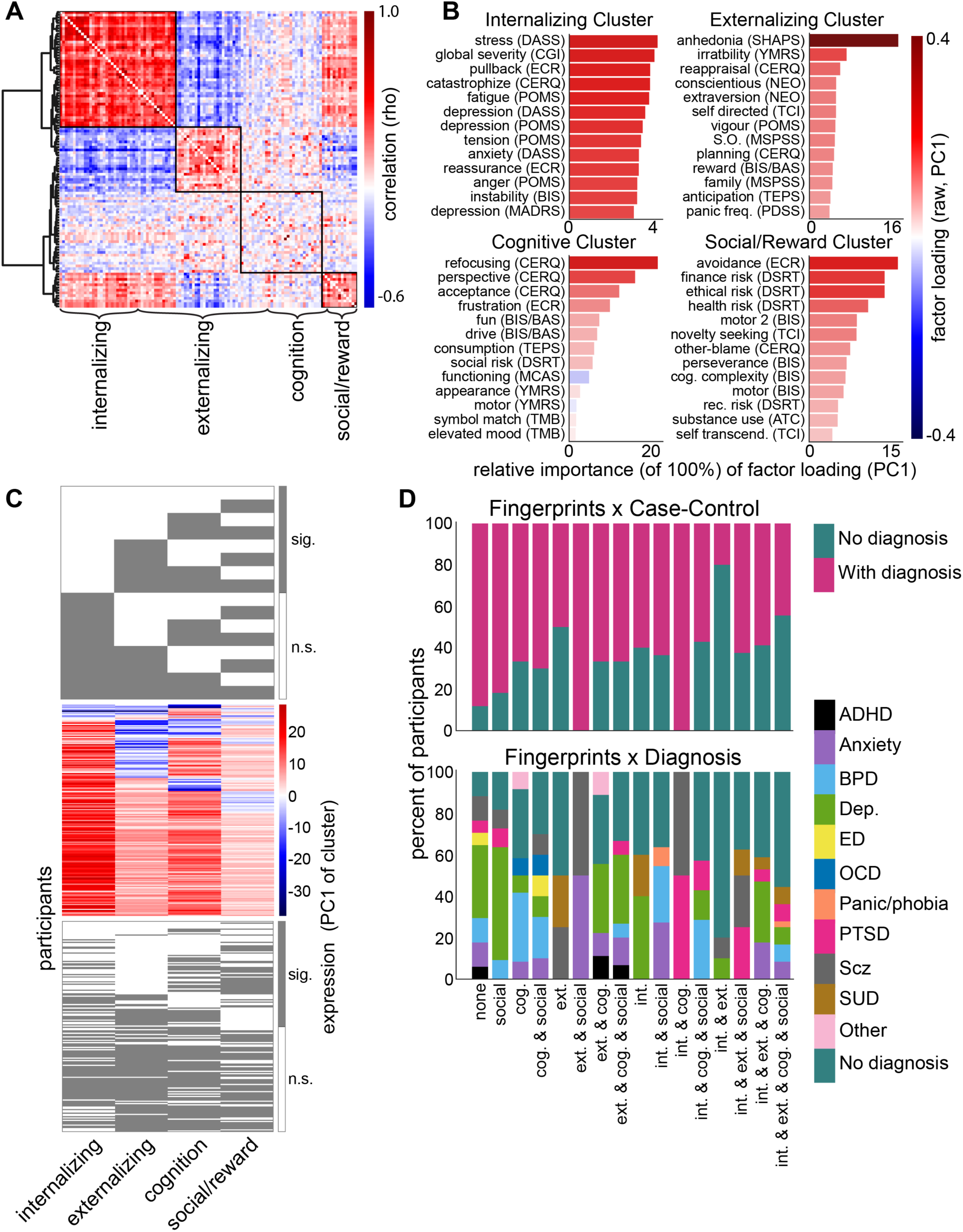
Uncovering dimensional symptom profiles. (**A**) Individual differences correlations (the extent that behavioral measure scores covary across individuals; rho) between all pairs of 110 behavioral measures (**see Methods; Supplemental Fig. S3**). Agglomerative hierarchical clustering was applied, and 18 metrics indicated a 4-cluster solution was optimal. Dendrogram: Ward’s distance. Black boxes outline clusters, named: internalizing, externalizing, cognition, and social/reward (see: **Supplemental Table S1**). (**B**) After PCA of measures in each cluster (independently assessed), the top measures (factor loadings) in PC 1 are shown (**Supplemental Fig. S4**); this guided naming conventions for the four dimensions of functioning in A and downstream. (**C**) Developing symptom fingerprints: each participant’s expression of the first PC of each cluster (**see Methods**). Top: all possible expression patterns (16 fingerprints), given by combinations of significant and non-significant expressions of each cluster. Middle: raw expression scores (log-likelihood) for all participants. Bottom: after testing whether each expression score in the middle panel was significant, they were binarized into one of 16 possible patterns per participant. (**D**) The percent-based distributions of case-control (top) and primary diagnosis (bottom) groups exhibiting each of the symptom profiles (or fingerprints) in C. Here, y-axes capture 100% of the total for each fingerprint individually. Abbreviations: int.: internalizing; ext.: externalizing; cog.: cognition; social: social/reward. Expected patterns relevant to quality assurance: participants diagnosed with depression were the largest proportion of the internalizing-only fingerprint; there was a diverse spread in the all-cluster (rightmost) fingerprint; and participants with anxiety diagnoses were in the externalizing and social fingerprint.

To quantify phenotypes that represented multi-dimensional information (i.e., all cluster expressions considered simultaneously) and were still differentiable, we developed a method termed *symptom fingerprinting*. With four clusters (dimensions of functioning), there were 16 possible combinations of binarized expressions (i.e., expressed cluster versus did not express cluster; **Fig. 3C**), which can also be thought of as symptom profiles. Here, expression was quantified via each participant’s score on the first PC of a given cluster (log likelihood; see **Methods**). One-sample t-testing was applied to clusterized expression scores and compared each participant’s score versus all other participants’ scores (multiple- comparisons corrected). Thus, binarization specifically refers to significant cluster expression versus non- significant expression (**Fig. 3C**). For quality assurance, we examined the distribution of participants’ primary psychiatric diagnoses and case-control statuses across the 16 symptom fingerprints (**Fig. 3D**). As expected, the fingerprint consisting solely of expressing the internalizing cluster included a dominant percentage of individuals with depression diagnoses. The fingerprint with all four clusters expressed had a diverse and more evenly spread distribution of individuals with and without psychiatric diagnoses. Perhaps reflecting the multiple domains of functioning impacted by PTSD^121,122^, individuals with that primary diagnosis were strongly represented in the internalizing-cognition fingerprint as well as the internalizing-externalizing-social fingerprint. Altogether we validated an empirically-driven approach to individually profile dimensional symptom structures that were exhibited by a transdiagnostic cohort of participants with and without psychiatric diagnosis.

### Brain network dynamics classify dimensional symptom fingerprints across health and disease

We used support vector classification (SVC^123^) to test the hypothesis that brain network dynamics are linked with dimensional phenotypes across health and psychiatric illness. Subgraph expression coefficients exhibited across six cognitive states (**Fig. 2**) were used to classify 16 symptom fingerprint labels (**Fig. 3**). To train the model, we randomly subsampled 80% of the discovery dataset (1000 permutations) and tested classification accuracy on the held-out validation dataset (see **Methods**). We built null models by randomly shuffling the fingerprint labels 1000 times to empirically obtain chance-level classification accuracy. Empirically-based chance accuracy was highly similar to theoretical chance, which was 1/16 or 6.25% (empirical chance=6.6%). When classifying fingerprints, there were two types of accuracy: strict and fuzzy. Strict accuracy was based on an all-or-nothing decision boundary of correct (100% accurate) or incorrect (0%). Given that symptom fingerprints were individualized profiles based on binarized *patterns* of cluster expression, we also considered a “fuzzier” decision boundary that accounted for the extent of overlap between predicted and actual fingerprint labels. Fuzzy accuracy was based on the percent of four possible cluster expressions that were correctly classified, and included possible values of 0%, 25%, 50%, 75%, and 100% on each permutation. An example fuzzy accuracy score of 50% is a predicted fingerprint label of internalizing-externalizing (1, 1, 0, 0) and an actual fingerprint label of internalizing-cognition (1, 0, 1, 0). This example model is half correct in identifying internalizing cluster expression, but half incorrect in mislabeling cognition-cluster expression as externalizing-cluster expression.

Symptom fingerprints were classified significantly above chance (strict: *t*(42)=7.2, *p*=3.8x10^-9^, *Cohen’s D*=17.8; fuzzy: *t*(42)=24.8, *p*=5.4x10^-27^, *Cohen’s D*=61.2; **Fig. 4A**). We implemented two comparison models with the same SVC pipeline except with the outcome labels of: (1) case-control status (*t*(42)=10.7, *p*=6.4x10^-14^, *Cohen’s D*=3.3), and (2) primary diagnosis group (*t*(42)=5.2, *p*=2.9x10^-4^, *Cohen’s D*=9.6; **Fig. 4A**). While brain network dynamics were able to significantly classify all approaches to grouping individuals, dimensional symptom profiles were classified with the largest effect size via both strict and fuzzy accuracy. This suggests that the boundaries between groups of individuals exhibiting symptom fingerprints are separated more precisely by brain network dynamics than case-control status or primary diagnosis categories.

**Figure 4.**
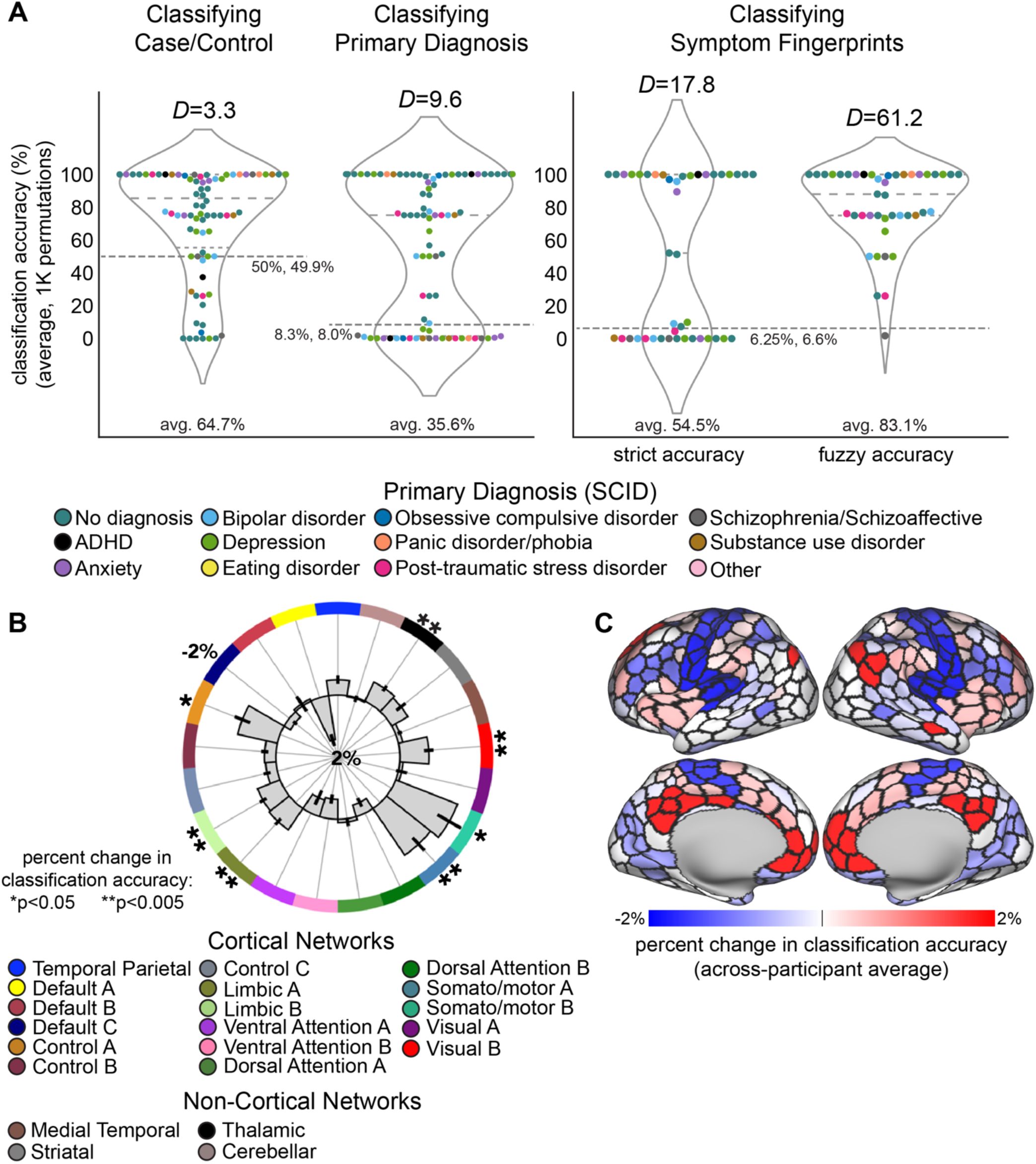
Brain network dynamics classified dimensional symptom fingerprints better than case-control status and primary diagnoses. (**A**) Classification accuracy of case-control status, primary diagnosis grouping, and symptom fingerprints (left-to-right) (dots: participants; colors, all plots: primary diagnosis) of the validation dataset. In each model, predictors included NMF-uncovered subgraph expression coefficients across all six cognitive states (three resting- and three task-states). Effect sizes (Cohen’s *D*) suggest that while all models were significant, classification accuracies of fingerprints (strict and fuzzy) were the most robust. *t*-statistics against empirical chance (dotted gray lines) based on 1000 permuted null models reported in-text. Theoretical and empirical chance (left, right) on dotted lines. (**B**) Classification of fingerprints was iteratively performed with each large-scale functional network withheld (simulated lesioning) before NMF. In each iteration, features underlying brain network dynamics did not include the lesioned networks’ FC estimates. A significant reduction in model performance suggests dynamics conferred by changing connectivity patterns in a given network were particularly important for classifying symptom fingerprints. Error bars: standard error of the mean across validation-set participants. (**C**) The same results as in B but projected onto a cortical surface (non-cortex indicated in B). Darker blue: functional brain networks whose across-state dynamics were particularly important for classifying dimensional symptom profiles.

Next, we examined the extent that each large-scale functional network (as in **Fig. 2A**) contributed to the accurate classification of individual symptom profiles. For each functional network, we iteratively applied simulated lesioning of FC estimates, within- and between-network. Then, we re-implemented NMF (**Fig. 2**) to uncover reconfiguration dynamics without brain regions from that network considered. Next, we classified symptom fingerprints (as in **Fig. 4A**, the “full model”) and quantified changes in accuracy from the full model. A statistically significant (nonparametric permutation testing; see **Methods**) decrease in accuracy indicated which connectivity patterns were influential in linking brain network dynamics with symptom profiles. These networks included: control (frontoparietal) A, limbic A/B, somato/motor A/B, visual B, and thalamus (**Fig. 4B-C**). This suggests that brain systems known to enable cognitive control processing (frontoparietal and thalamus) and networks linked with task-relevant resource processing (visual and somato/motor) exhibit dynamics that are prominently linked with dimensionally-organized cognitive and behavioral functioning across health and psychiatric disease.

### Cognitive control processing influences the link between brain network dynamics and dimensional symptom profiles

To test how influential each cognitive state was to the link between brain network dynamics and symptom fingerprints, we implemented NMF with varied combinations of input connectivity patterns. This allowed us to differentiate context-specific shifts in connectivity from general rest-to-task shifts in connectivity (as in **Fig. 4**). The seven comparison models included these inputs to NMF: (1) all states (**Fig. 4**; a general reference model); (2) resting-state FC only, which likely contains small shifts in intrinsic^18,31,105,124^ network organization (not task-linked); (3) task-state FC only, consisting of Stroop^96^ and Emotional Faces^125^ FC; (4) resting- and Stroop-task-state FC, likely containing shifts in intrinsic connectivity relevant to cognitive control processing; (5) resting- and Emotional Faces-task-state FC, with shifts relevant to processing fearful faces; (6) Stroop FC only, containing static FC patterns relevant for cognitive control; and (7) Emotional Faces FC only, containing static FC patterns relevant for fear processing.

Across all models, symptom fingerprints were classified with greater effect size than case-control status and primary diagnostic group (**Fig. 5A-D**). As expected, there was a broadly consistent pattern of model performances between strict-accuracy and fuzzy-accuracy (**Fig. 5C-D**). The all-states model exhibited the lowest performance (strict accuracy: *D*=17.25; fuzzy accuracy: *D*=54.73), suggesting that network dynamics across wider-spanning task contexts may yield suboptimal decision boundaries separating clinical symptom structures expressed across individuals. The rest-and-Stroop model performed the best (strict accuracy: *D*=18.85; fuzzy accuracy: *D*=62.91), suggesting that connectivity shifts between an intrinsic state and a cognitive control state accounted for individual clinical variability particularly well. Interestingly, case-control and primary diagnosis were classified with a similar pattern across models. The main difference was that all-states performed worst and best in classifying primary diagnosis and case-control statuses, respectively. These data suggest that separability of case-control status may be sensitive to the number of cognitive states (or, amount of data) used to uncover network dynamics. Primary diagnoses may be classified best by dynamics across similar cognitive contexts. Conversely, dimensional symptom profiles may be separated best by a specific, foundational shift in cognitive context.

**Figure 5.**
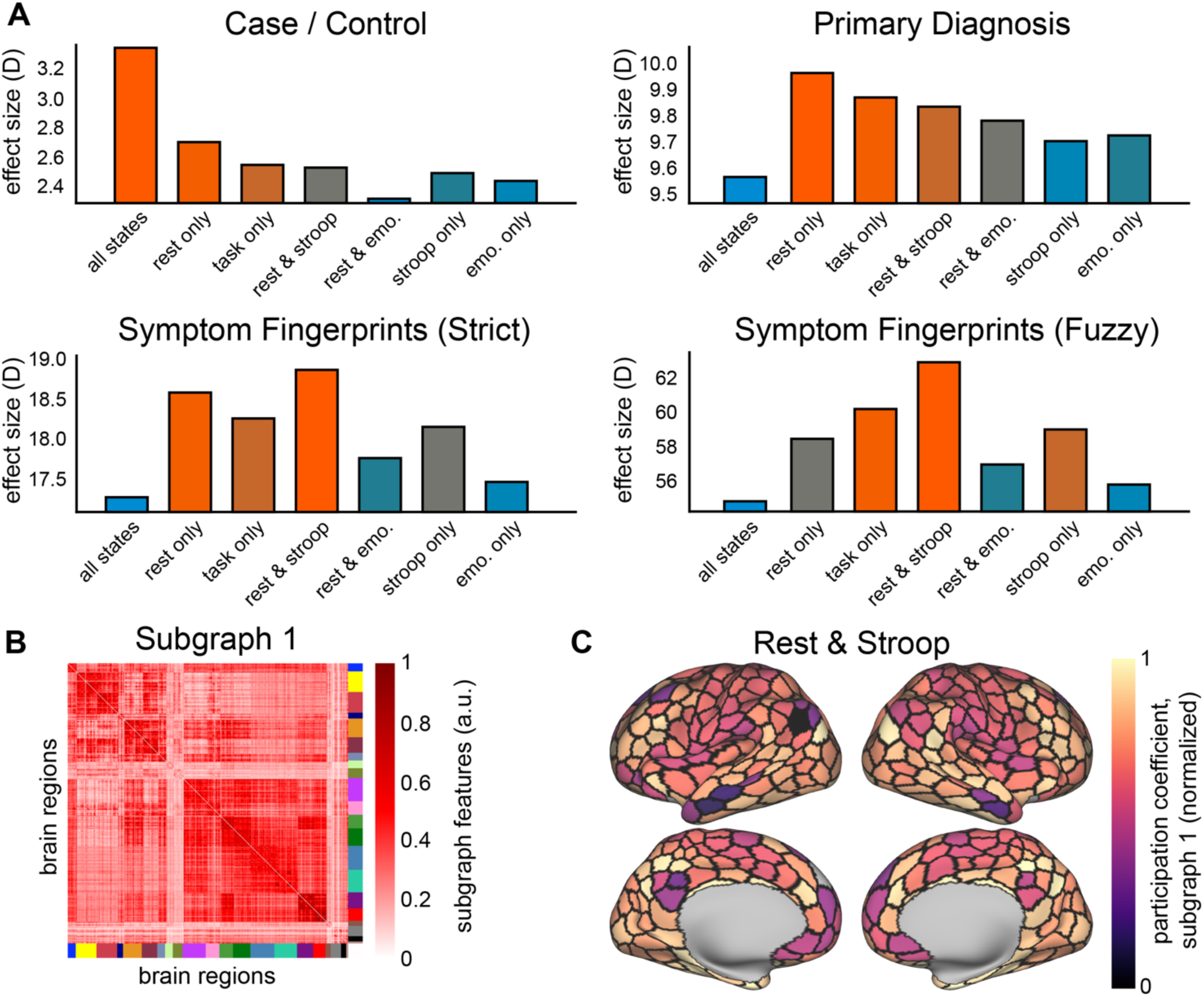
Reconfiguring connectivity patterns from rest-to-Stroop task states prominently influence the classification of dimensional symptom profiles. (**A**) Classification models with connectivity patterns from different cognitive states inputted to NMF (model inputs indicated on x-axes; see **Methods** and text for details). Headers indicate model comparison results for classifying case/control status, primary diagnosis grouping, symptom fingerprints with strict accuracy, and fingerprints with fuzzy accuracy, respectively (effect size of cross-participant model accuracies given by Cohen’s *D*). Bars are color-coded from blue-to-orange to indicate worst-to-best model performance. In the models assessing how well brain network dynamics classify dimensionally-organized symptom fingerprints, reconfigurations across rest-to-Stroop performed best, suggesting that information processing shifts from an intrinsic state to a cognitive control state account best for the expression of symptom profiles. (**B**) NMF-uncovered subgraph 1, reformatted into whole-brain region-by-region networks. This was used in (**C**): regional participation coefficient (normalized) of the best performing model (rest & Stroop) for classifying fingerprints (**Supplemental Fig. S5 for other models**).

Further, we found that control (frontoparietal) network regions of the first NMF-uncovered subgraph (**Fig. 5E**) in the rest-and-Stroop model strongly exhibited a functional hub property based on participation coefficient^126^ (**Fig. 5F; Supplemental Fig. S5**). Additionally, striatal, thalamic, and default C networks included strong hub regions and the default A/B, somato/motor A/B, dorsal attention B, and salience/ventral attention A networks exhibited the fewest hubs. For the majority of networks, hubs either stayed consistent or slightly decreased in the rest-and-Stroop model versus the rest-only model, suggesting that the increased prediction accuracy in the rest-and-Stroop model (**Fig. 5C-D**) was not solely driven by increased diversity in network interaction patterns while engaging with a cognitively demanding task. We propose that accurate separation of symptom fingerprints is influenced by the dynamic shift in cognitive context from an intrinsic state to a more cognitively demanding state. Altogether, and considering **Fig. 4** results uncovering the importance of networks linked with cognitive control, we propose that the flexible recruitment of cognitive resources to meet increased task demands is a key determinant of dimensional symptom structures exhibited across health and psychiatric disease.

## Discussion

We provide evidence that brain network dynamics are linked with transdiagnostic symptom profiles across health and disease (**Fig. 1**). The application of NMF to whole-brain functional connectivity patterns across varied cognitive states (**Fig. 2**) allowed us to examine dynamic constraints upon across-state shifts in functional network interactions^86^. Individuals with psychiatric diagnoses exhibited flattened (i.e., less differentiated) network dynamics, particularly across resting- and Stroop-task states. These data suggest that cognitive control/executive functioning is disrupted across psychiatric illnesses^45–49^, and that suboptimal recruitment of processing resources may emerge via flattened network dynamics. Hierarchically-informed methods^75–80^ were used to characterize symptom fingerprints, or individual symptom structure profiles that mapped to core dimensions of functioning (**Fig. 3**). Four well-established dimensions of functioning were evident across participants: internalizing, externalizing, cognitive, and social/reward (**Supplemental Table S1**^110^). Symptom fingerprints – which profile the extent that an individual expressed each of these dimensions – were accurately classified by brain network dynamics. When compared to primary diagnostic group and case-control status, symptom fingerprints exhibited the largest effect size (**Fig. 4**). Simulated lesioning of functional brain networks revealed that cognitive control-linked systems (frontoparietal and thalamus) and task-relevant resource systems (visual and somato/motor) were most important to the link between brain network dynamics and symptom fingerprints. Reconfiguration dynamics across resting-to-Stroop states separated fingerprints with the highest accuracy (**Fig. 5**), further corroborating the clinical relevance of cognitive control processing. Altogether, these results reveal that individualized symptom profiles exhibited across four core dimensions of functioning can be accurately accounted for by brain network dynamics.

Resting-state dynamic FC has been shown to outperform static FC and composite static-dynamic FC models in classifying schizophrenia, bipolar disorder, and healthy controls^61^. In the present study, we examined a transdiagnostic sample that consisted of 11 primary psychiatric diagnosis groups and investigated the extent that whole-brain network dynamics exhibited across rest- and task-based cognitive states can account for varied models of health and disease. We demonstrated that brain network dynamics can discriminate between dimensionally-organized symptom profiles, case-control status, and primary diagnoses with high accuracy. This supports the proposition that brain dynamics contain more process-relevant information than static network features and thus confer an enhanced separation of the boundaries between diagnostic groups (**Fig. 1**). This is consistent with recent evidence suggesting that ADHD diagnoses are accounted for by increased range but decreased fluidity in dynamic FC, which was not discoverable via static FC^58^. Similarly, individuals with schizophrenia exhibit: altered dwell times across varied states^53^; transient connectivity profiles predicting active psychosis^52^; shorter dwell times in states engaging the frontoparietal network^54^; alterations to dynamic shifting between brain states^55^; and less complex trajectories through connectivity state-space^56^. Dysfunctions in information processing across brain systems have also been linked with less dynamic connectivity repertoires^127,128^, deviations from healthy reconfiguration statistics^38^, and altered microstate topography in select EEG frequency bands^129^.

The present study provides key insights on the extent that across-state network reconfigurations track dimensions of cognitive and behavioral functioning. However, *why* brain network dynamics account for behavior better than static network features remains an open question. We provide evidence that brain connectivity patterns reconfiguring in response to shifts in task context are strongly linked with core domains of functioning expressed across all participants (with and without diagnoses). Building on this (as well as key findings in the literature^51,52,61,70,127,130^), it is likely that multi-scale, integrated mechanisms of information processing^99,131^ are embedded in, and revealed through, brain network dynamics, which are in turn approximated by static network properties^3,132^. One framework that is particularly relevant to psychiatry is that, while the brain is a dynamical system, so too is symptom expression^4,133^, as many illnesses involve cycling or phases over time. In both neural systems and psychiatric symptomatology, research has characterized dynamical regimes with patterns of dominating (attractor) and less stable (repeller) states. For example, obsessive compulsive disorder has been linked with over-stability of attractor states, recapitulated across brain systems at multiple scales^57^. Moreover, dynamical systems theory has begun to infer the neurobiological relevance of transitioning between such states^17,134^, as well as the energetic costs linked with the processing resources needed for different functions^135,136^. In this account, network dynamics are more closely aligned to dimensionally-organized axes of psychiatry than static network properties (**Fig. 1**). This is a key hypothesis that future work should address by building on the present methods with longitudinal data collected synchronous to symptom expression. A related framework posits an intrinsic (baseline) set of statistics that characterize brain network dynamics that give rise to task-induced modifications^137–140^. Here, the controllability of select brain regions is quantified^138^, and shifts in network dynamics may be associated with altered information entropy and metabolic costs^141^. This framing is not directly competitive with dynamical systems theory but does make nonequivalent predictions. For example, a key hypothesis would be the extent that network dynamics account for symptom structure is determined by a breakdown in the efficiency of information processing across varied task contexts.

There are important considerations when interpreting standard dynamic FC estimates, such as sliding window correlations, particularly with respect to non-stationarity^132^. Time-invariant FC assumes that the statistics underlying network interaction patterns are static or fixed over time. Therefore, the careful implementation of dynamic FC may improve neurobiological plausibility by accounting for non-stationary statistical structure in neural processes. However, a critical confound in standard approaches to dynamic connectivity is that such statistics are also influenced by non-neural, nuisance sources^142^. This is particularly problematic with psychiatric participants, given increased motion artifacts^143,144^, respiratory effects^145,146^, and altered arousal processing^147,148^. We sought to address these concerns in three ways. First, the present data were processed with careful consideration of motion correction, denoising, and quality control benchmarking^110^. Second, we used a train-test-validation data splitting scheme to account for data leakage and reduce spurious effects^149,150^ (**Supplemental Fig. S6**). Third, we used a method for uncovering constraints upon brain network dynamics that depended chiefly on the structure of the input data, and contained no free parameters linked with events or windows of time. To make non-stationarity inferences, we instead opted for a model comparison approach (**Fig. 5**), where connectivity patterns relevant to varied cognitive states were considered to compare reconfiguring network interactions across resting-state-alone, rest-to-Stroop, task-alone, and so forth. The present study advances clinical neuroscience by bypassing undue influence from non-neural sources of variance, however, we encourage future work that systematically compares how well other brain network dynamics methods can discriminate the structure of psychiatric symptoms.

Finally, cognitive control processing was prominent in establishing a robust link between network dynamics and symptom fingerprints (**Figs. 2, 4-5**). This is consistent with evidence that the flexible recruitment of processing resources is negatively impacted across psychiatric diagnoses^16,45,49,151^. However, the extent that network dynamics are altered across cognitive control systems in psychiatric patients – as well as the direction, stability, and generalizability of this effect – remains an important empirical question for future research. The current approach can be readily expanded on. For example, we used state-general estimates of FC – that is, connectivity estimates for the entire duration of the Stroop task, Emotional Faces task, and resting-state. This could be refined in future work by estimating FC based on task context, condition, or other temporally-linked factors. Relatedly, connectivity can be estimated by alternatives to product-moment correlations, such as the improved validity given by regularized regression^100^. NMF has also been successful with multivariate data inputs, such as structural and functional connectivity^88^, which may be an important extension of the present work given evidence that properties of the structural connectome constrain the spread of psychopathological brain processes^130^. Lastly, we are enthusiastic about future work that expands upon transdiagnostic symptom fingerprinting, for example, probing if the extent of differentiation in brain network dynamics scales with the extent that dimensions of functioning are expressed in individual symptom profiles.

Here, we studied a transdiagnostic sample of participants with and without psychiatric diagnoses and demonstrated that whole-brain network reconfiguration dynamics can discriminate symptom fingerprints with high accuracy. Cognitive control processes and associated brain networks were prominently implicated in these data, suggesting that executive dysfunction is key to developing a brain-process-level account of transdiagnostic psychopathology. Lastly, this work benchmarks a readily adaptable pipeline for investigating brain-behavior links with improved neurobiological plausibility (**Fig. 1**) as well as accounting for known limitations in using discrete diagnostic boundaries in clinical neuroscience.

### Online Methods *Data collection* Participants

Data were collected as part of the Transdiagnostic Connectome Project (TCP)^110^ (**Supplemental Fig. S6**) at the FAS Brain Imaging Center at Yale University in New Haven, Connecticut, US, and the Brain Imaging Center at McLean Hospital in Belmont, Massachusetts, US. All participants were given written informed consent documentation following the protocols of each center’s Institutional Review Board. Full details of the TCP dataset can be found elsewhere^110^, but in brief, N=241 participants were 18 through 70 years old (mean=36.52, SD=13.09 years) and were recruited from the community as well as patient referrals from collaborating psychiatric clinicians. Participants were screened to assess if they met the following inclusion criteria: that they were eligible for MRI scanning, were not colorblind, and were not previously diagnosed with a neurological disorder or abnormality. To maximize data available for functional network analyses in the present study, N=219 TCP participants were included given that these 219 participants had neuroimaging data for six of the seven fMRI runs. Of this N=219 in the present study (mean age=36.2, SD=12.9 years; n=93 (42.5%) identified as male, n=123 (56.2%) as female, and n=3 (1.4%) as nonbinary), there were n=134 (61.2%) with psychiatric diagnoses, n=85 (38.8%) without a history of psychiatric diagnoses or treatment (**Supplemental Fig. S6C**; and see sections below for further details on MRI data acquisition and brain network dynamics analyses).

### Study outline

After initial screening, there were three study sessions in the TCP data acquisition pipeline (**Supplemental Fig. S6A**). First, an in-person session that consisted of demographic and health surveys, the Structured Clinical Interview for DSM-5 (SCID-V-RV^104^), a battery of clinician-administered scales, and self-report scales. Second, an in-person session that included MRI scanning (see *MRI data acquisition* section below for further neuroimaging details) as well as further self-report scales. Third, an online session of supplemental self-report scales as well as the TestMyBrain suite^117^. Scales and surveys included a variety of cognitive, behavioral, and clinically-relevant measures and will be collectively referred to as “*behavioral measures”* in the present study (see *Dimensional symptom analysis* section below for further details).

Based on the SCID-V-RV, eleven primary psychiatric diagnosis groups were present (**Supplemental Fig. S6**) including attention-deficit/hyperactivity disorder (ADHD), anxiety, bipolar disorder (BPD), depression, eating disorder, obsessive compulsive disorder (OCD), panic disorder/phobia, posttraumatic stress disorder (PTSD), schizophrenia/schizoaffective, substance use disorder, and other. Select diagnostic groups were concatenated based on central symptomatology, as follows: anxiety refers to any anxiety- central diagnosis such as generalized anxiety disorder, social anxiety, and anxiety not otherwise specified (NOS); BPD included BPD I, BPD II, and cyclothymia; depression included major depressive disorder, dysthymia, and depression NOS; SUD included addiction to any substance and/or alcohol (e.g., cocaine use disorder and alcohol use disorder combined into one SUD category); and other refers to rare diagnoses such as premenstrual dysphoric disorder and psychosis NOS. Note that in downstream analyses, there were 12 labels for primary diagnosis; the 12th label was “no diagnosis”, referring to the healthy comparison participants without psychiatric diagnoses based on the SCID-V-RV.

### Data splitting scheme

It was possible that in the final analysis testing the extent that functional brain network dynamics can classify symptom structure (described in following sections) there would be leakage of information between training and test participants^149,150^. This was because prior analytic steps required either supervised learning as well (as in random forest imputation; see *Dimensional symptom analysis* section below) or hyperparameter optimization with cross validation (as in nonnegative matrix factorization; see *Brain network dynamics analysis* section below). This potential for leakage between participants has been found to artificially inflate prediction results, which can be problematic for interpreting neurobiological links with behavior. Note that data collection site, familial relations, and age were not found to be problematic with respect to data leakage impacting modeling results.

To mitigate leakage across participants, we implemented a data splitting scheme at the outset of the present study (**Supplemental Fig. S6B-C**). Of the N=219 total participants, approximately 80% (n=176) were allocated to a discovery dataset and approximately 20% (n=43) to a held-out validation dataset, which was left untouched until the final analyses. The discovery set was randomly split further for various analysis steps prior to classification into training and test sets. To ensure that the underlying structure of the total dataset was captured by each of the discovery and validation sets, two percentage distributions were constrained: (1) proportion of participants in each primary diagnosis category, and (2) proportion of participants from each collection site (**Supplemental Fig. S6C**); otherwise, participants were allocated randomly.

### MRI data acquisition

All MRI data were collected at either the Yale University FAS Brain Imaging Center or the McLean Hospital Brain Imaging Center. The best practices put forth by the Human Connectome Project (HCP)^152,153^ were followed as closely as possible in both MRI acquisition protocols and data processing. Both scanners were Siemens Magnetom 3T Prisma models with 64-channel head coils. Whole-brain and multi-echo MPRAGE sequences were used to collect T1w anatomical data with the following specifications: repetition time (TR)=2.2 s; echo times (TE)=1.5, 3.4, 5.2, and 7.0 ms (root mean square of each echo used to compute a single image); flip angle=7°; inversion time=1.1 s; sagittal orientation for phase encoding=anterior (A) to posterior (P); slice thickness=1.2 mm; total slices acquired=144; resolution=1.2 mm^3^. Specifications unique to T2w anatomical images were: TR=2.8 s; TE=326 ms. Whole-brain, multiband, and echoplanar (EPI) functional MRI data were acquired with the following specifications: TR=0.8 s; TE=37 ms; flip angle=52°; voxel size resolution=2 mm^3^; multiband acceleration factor=8. There were three cognitive states assessed during scanning plus variants based on AP/PA phase encoding direction, totaling 7 fMRI runs altogether: (1) 2 resting state fMRI runs in both AP and PA (4 total); (2) Stroop inhibitory control task^95,96^ fMRI in both AP and PA (**Supplemental Fig. S6D**) (2 total); and (3) Emotional Faces task^125^ fMRI in the AP direction (**Supplemental Fig. S6E**). There were 488 volumes (TRs) collected for each resting-state fMRI run; 510 TRs for each Stroop task fMRI run; and 493 TRs for Emotional Faces task fMRI. See [^110^] for further details on TCP MRI data acquisition protocols as well as full details on paradigms used during task fMRI (**Supplemental Fig. S6D-E**). As previously noted, only six of the seven TCP fMRI runs were included in the present study to maximize available data for N=219 participants: resting-state run two, PA, was excluded, leaving a balanced total of three resting- and three task-state fMRI runs for downstream analyses.

### MRI data processing

The HCP minimal processing pipelines^152^, version 4.7.0 were applied to neuroimaging data. HCP best practices, technical specifications, and quality assurance analyses are detailed extensively elsewhere^110,152^, but briefly we implemented: anatomic reconstruction and segmentation; motion correction; EPI reconstruction, segmentation, and spatial normalization to a standard template; intensity normalization; multimodal surface matching (“MSM-All”) registration^154^; single-run ICA-FIX denoising^155,156^; and de-drift and resampling. Resulting data were in CIFTI 91k-vertex grayordinate space, which were then parcellated (i.e., average time-series of enclosed vertices) into 400 cortical regions^106^, 32 subcortical regions^107^, and two cerebellar regions^108^, totaling 434 whole-brain regions. Note that in all figures and text, the term *region* refers to either a functionally parcellated area of cortex or an anatomical subdivision of subcortex or cerebellum. The term parcel or node is sometimes used for cortical brain areas, but we adopted the general term *region* for subdivisions of both cortex and non-cortex for simplicity. In both cortex and non-cortex, a brain *network* refers to a functionally- or anatomically-linked collection of regions (see **Fig. 2** in Results for parcellated regions projected onto a standard brain surface and volume for cortex and non-cortex, respectively; color-coded by network assignment).

### Functional connectivity estimation and functional network partition

Functional connectivity (FC) was estimated for each participant and functional scan via product-moment correlation between each pair of regions’ minimally processed and denoised timeseries. These 434 region by 434 region FC matrices were further partitioned into large-scale functional networks based on a widely-used 17 cortical network solution^106,124^, three subcortical networks^107^, and one cerebellar network^108^; totaling 21 whole-brain functional networks (see **Fig. 5** in Results). Fisher’s Z-transformation was used whenever FC estimates were averaged.

### Brain network dynamics analysis

Following prior work^86,88,90–92^, we implemented an unsupervised machine learning tool called nonnegative matrix factorization (NMF) to uncover hidden constraints upon functional brain network dynamics. NMF is known for being a parts-based decomposition of multi-dimensional input data (matrices or tensors) into two lower ranking matrices; one with features that approximate a basis set (also termed motifs, weights, or subgraphs), and another being an encoding matrix of coefficients (also termed expressions or expression coefficients). Standardly, all matrices and/or tensors are non-negative. NMF-uncovered features are optimized for a linear approximation of the original input data and are not constrained by independence or orthogonality (as in principal components analysis), which is likely more neurobiologically appropriate because flexibly co-occurring and/or overlapping network structures are permitted.

In the present application, FC estimates (upper triangle and positively-thresholded values) for each participant and fMRI scan were stacked vertically (as in [^86^]), and this network configuration matrix was used as the input to NMF. In this usage, across-state reconfiguration dynamics are captured by the presence of both resting-state and task-state FC estimates. Here, NMF is thought to uncover features of functional brain network interactions that constrain dynamic transitions across varied cognitive states. While unsupervised, NMF requires three parameters to be optimized: k, or the number of feature matrices; alpha, or the multiplicative regularization term for factorized matrices; and beta, which is a loss parameter based on minimizing divergence. We cross-validated the discovery dataset by splitting participants randomly and evenly into training and test sets over 100 folds (note that this was limited by computational feasibility) to tune these parameters, which were then applied to NMF of the full discovery dataset as well as the validation dataset. Possible values of k ranged from two through 20 (steps of one); alpha ranged from 0.1 to 2.0 (steps of 0.1); and beta ranged from 0.2 to 2.0 (steps of 0.2). Accuracy of the cross validation was assessed via reconstruction error^86,157^ as well as the nonparametric Mantel correlation^158^ with the actual FC configuration matrix. To implement NMF, we used the Python toolkit provided by scikit-learn (version 1.6.1) with the multiplicative update solver, and all other parameters were set to their defaults. Accuracy was stable with the following parameter values: k = 5, alpha = 0.2, beta = 1.2 (see Results **Fig. 2** for NMF pipeline schematic; **Supplemental Fig. S1**).

### Dimensional symptom analysis

The TCP dataset consists of a rich variety of 163 behavioral, cognitive, and clinical measures (here, collectively referred to as *behavioral measures* for simplicity; see **Supplemental Table S1** table for full list of all clinical, self-report surveys, and cognitive batteries used herein), which we used to uncover dimensionally-organized symptom structures^75–81,111^, that we termed symptom *fingerprints* or *profiles*.

First, we curated the data to exclude redundancies. One type of redundancy was the presence of a summary score and subscale scores; here we prioritized subscale scores to cover the most amount of behavioral variance unless the distribution of subscale scores were highly inflated at a single point or otherwise highly non-normal. Another type of redundancy was the presence of both a raw score and a t-stat; here we followed prior literature for content-specific best practices. Then, select variables were excluded for theoretical reasons, such as the childhood trauma questionnaire^159^, which is an assessment of predisposing psychiatric risk factors based on traumatic experiences in childhood, while all other measures were referential to current state or recent past (i.e., adulthood). The final set of measures totaled 110 clinical, behavioral, and cognitive variables.

After curation, the distribution of each behavioral measure was assessed for Gaussianity with four metrics: D’Agostino’s K-squared test, the Shapiro-Wilk test, the Anderson-Darling test, and Kolmogorov-Smirnov test. If a given measure exhibited a non-normal distribution based on more than two of these metrics, it was submitted to the bestNormalize package (version 1.9.1) in R^160^, which samples a suite of transformations and suggests the most consistent and accurate function for normalizing the given data (see **Supplemental Fig. S3** for representative examples). After this initial transformation, the four Gaussianity metrics were implemented again to identify measures that were still highly non-normal. Here, the remaining variables were all zero-inflated, which we opted to binarize for interpretability.

Following normalizing and binarizing transformations, select behavioral measures were unavailable in some participants. To account for this, we used random forest imputation (also termed “miss forest”)^161^ based on prior evidence that it is robustly applicable to nonparametric and/or mixed data types. Random forest imputation is a supervised learning algorithm that we performed on the discovery dataset; where 80% of participants were randomly allocated to the training set and 20% to the testing set. In order to ensure that the variety of native score ranges did not impact downstream analyses, resulting data (i.e., behavioral measures that were transformed and imputed) were scaled to the same space with min-max normalization between the values of zero and one. This approach is widely used in machine learning and has the benefit of recapitulating the original shape of the distribution, just shifted between a fixed minimum and maximum value.

Following prior work, we applied agglomerative hierarchical clustering with Ward’s linkage distance^162^ to individual differences correlations (here, Spearman’s rho) of all pairs of behavioral measures. An individual differences correlation estimates the extent that scores on a given pair of behavioral measures covaries across individuals, or how well individual differences are matched in each measure-to-measure pair. If individual differences are highly correlated for a given pair of measures, then they are capturing a similar dimension of functioning and will be allocated to the same cluster. Throughout the present study, the term *dimensionally-organized* refers to this clusterization. The optimal number of clusters (here: 4 clusters) was found with the R package NbClust (version 3.0.1)^163^, which uses 30 performance metrics to determine the best clustering solution. Then, we implemented principal components analysis (PCA) on each of the four clusters of variables (i.e., the original scores, not correlation values), and used the first PC for downstream analyses. This approach has the benefit of similar amounts of variance explained being represented for the first PC of each cluster (see **Supplemental Fig. S4** and **Table S1**), as well as including each behavioral measure in only one cluster to prevent overfitting to any given measure (which may happen if PCA were performed directly without the hierarchical clustering step). Per participant, we used an output from scikit-learn called score_samples to estimate the extent (given by log-likelihood) that each participant expressed a given cluster. Following prior work^80,112,113^, clusters were named for interpretability, however we caution against overly interpreting this naming system (as well as priorly used naming systems). To partially account for potential researcher bias in naming clusters, one researcher pre-labeled each of the 110 behavioral measures with broad dimensions of cognitive or behavioral functioning (referred to as dimensions of functioning in the main text), which was corroborated by two other researchers (**Supplemental Table S1**). Then, following clustering and PCA, the percentages of each label in each cluster – weighted by normalized factor loadings – were summarized programmatically. This guided naming the four clusters as follows: internalizing, externalizing, cognition, and social/reward.

Lastly, individual cluster expression scores were converted into categorical patterns or profiles, termed *symptom fingerprints*. For each participant, we performed one sample t-tests to test the null hypothesis that a given cluster expression score was equal to all other participants’ scores (alternative: participant’s score was greater than all other scores). We corrected for multiple comparisons with the false discovery rate^164^. A pattern of zeros and ones were assigned for significant and nonsignificant cluster expression (per participant), respectively. This binarization pattern allowed for 16 possible combinations (or *fingerprints*) of cluster expressions (see Results **Fig. 3**) which were used as 16 possible symptom fingerprint labels in downstream analyses.

### Classification analysis

In order to test the extent that functional brain network dynamics link with dimensional symptom structures across a transdiagnostic and non-diagnosed sample, we used nonlinear support vector classification (SVC) by implementing the Python toolkit NuSVC provided by scikit-learn^123^. NMF-uncovered subgraph expression coefficients across each of the six states (per participant) were used as the predictors and symptom fingerprints (16 possible), primary diagnosis (12 possible), and case-control (2 possible) labels were used as to-be-classified labels in three separate models, which were later compared. Over 1000 permutations, 80% of discovery set participants were randomly allocated for training, and the held-out validation set was used for testing. Accuracy was based on an average of these permutations. Each model had a different theoretical chance level: fingerprints: 6.25%; primary diagnosis: 8.33%; case-control: 50%. These were each empirically corroborated by nonparametric permutation testing, where 1000 null models were built on randomly shuffling participant labels^165^. Note that empirical chance closely matched theoretical chance in each model (see **Results** for full reporting). Statistical testing was performed on accuracy values with one sample t-tests against empirical chance, and Cohen’s *D* effect sizes were used to compare the three models. An added benefit of the fingerprinting approach was that it allowed us to develop a nuanced accuracy metric, which we termed “fuzzy” accuracy (and henceforth referred to the original as “strict” accuracy). Fuzzy accuracy accounted for overlap in fingerprint patterns, for example: label one was based on significant expression of all four clusters (1, 1, 1, 1) and label two on three of the four clusters (1, 1, 1, 0); a misclassification between labels one and two was considered 75% correct in terms of fuzzy accuracy and 0% correct in terms of strict accuracy.

### Model comparison

We performed two types of model comparison where connectivity patterns inputted to NMF were modified, NMF was re-implemented, and classification of case/control status, primary diagnosis group, and symptom fingerprints were re-assessed based on the modified model of brain network dynamics. Following this, effect size (Cohen’s *D*) of cross-participant model accuracies (in all models, accuracy given by averaging 1000 permutations) were compared. First, we used this approach to simulate lesioning of functional network brain regions and compare the relative importance of brain systems to downstream classification. Each set of regions was withheld from NMF one network at a time at a time (i.e., models were iterated over 22 networks). Model performance before and after lesioning was compared and any statistically significant reduction in model accuracy was interpreted as that functional system being important for separating class labels. Second, we used a similar approach to compare the relative importance of connectivity estimates yielded by different sets of cognitive states (but here, all brain regions were included in all models). The following models were tested: (1) all states: the original or “full” reference model with FC inputted to NMF from all six cognitive states; (2) rest only: NMF uncovered dynamic shifts in connectivity patterns across resting-state (i.e., intrinsic processing) only; (3) task only: shifts across inhibitory cognitive control (Stroop) to emotional faces task states; (4) rest & Stroop: shifts from intrinsic states to cognitive control states; (5) rest & emotional faces: shifts from intrinsic states to emotional-faces recognition states; (6) Stroop only: shifts across cognitive control task states only; and (7) emotional faces only: shifts across emotional faces task states only. Models 2, 6, and 7 were not expected to include large shifts in information processing or cognitive context (i.e., underlying dynamics are likely stable), whereas models 1, 3, 4, and 5 parameterized varied shifts in cognitive context.

## Data availability

All analysis code is openly available here: https://github.com/HolmesLab/ClinicalNetDynamics, which was primarily written in Python version 3.12.1. TCP unprocessed and processed neuroimaging data as well as all behavioral measures are openly available^110^ here: https://nda.nih.gov/study.html?id=2932 and here: https://openneuro.org/datasets/ds005237.

## Acknowledgements

We would like to sincerely thank all of the participants for their significant contribution to this research. This work was supported by the National Institute of Mental Health (R01MH123245 to AJH and R01MH120080 to AJH). All conclusions, inferences, opinions, findings, suggestions, and recommendations expressed or otherwise presented in this manuscript are those of the authors and do not reflect the views of the funding bodies. The authors acknowledge the Office of Advanced Research Computing at Rutgers University as well as the Yale Center for Research Computing at Yale University for providing access to high performance compute clusters and associated resources essential to this research.

