## Supplemental for "Brain network dynamics reflect psychiatric illness status and transdiagnostic symptom profiles across health and disease"

### Supplementary Information

This supplemental material includes *Supplementary Tables*, *Supplementary Figures*, and *Supplementary References*.

### Supplementary Tables

**Table S1. 110 behavioral, cognitive, and clinical measures used for symptom fingerprinting.**

| Measure name (full) | A priori labels | Cluster assignment | Factor loading (rel. %) |
|---|---|---|---|
| Positive and negative syndrome scale (negative subscale) | internalizing, thought disorder, global | internalizing | 0.4 |
| Positive and negative syndrome scale (positive subscale) | thought disorder, global | internalizing | 0.5 |
| Broad autism phenotype questionnaire (rigid subscale, avg.) | social, externalizing, global | internalizing | 1.1 |
| Broad autism phenotype questionnaire (aloof subscale, avg.) | social, externalizing, global | internalizing | 1.1 |
| Positive and negative syndrome scale (general subscale) | global, thought disorder | internalizing | 1.1 |
| Range of impaired functioning tool (total) | social, externalizing | internalizing | 1.2 |
| Barratt impulsiveness scale (subscale 1, self control) | externalizing, healthy functioning | internalizing | 1.2 |
| Barratt impulsiveness scale (subscale 2, non-planning) | externalizing, healthy functioning | internalizing | 1.6 |
| Broad autism phenotype questionnaire (pragmatic language, avg.) | social, externalizing, global | internalizing | 1.6 |
| Retrospective self-report of inhibition (social/school subscale) | internalizing, externalizing | internalizing | 1.6 |
| Profile of mood states (confusion subscale t-score) | internalizing, thought disorder | internalizing | 2.0 |
| Positive urgency measure (total, avg.) | externalizing | internalizing | 2.0 |
| Experiences in close relationships inventory (avoidance subscale) | social, externalizing | internalizing | 2.0 |
| Cognitive failures questionnaire (forgetfulness subscale) | global, thought disorder | internalizing | 2.1 |
| Behavioral inhibition/activation scale (inhibition subscale) | reward, internalizing | internalizing | 2.1 |
| Anxiety sensitivity index | internalizing | internalizing | 2.1 |
| Cognitive failures questionnaire (false triggering subscale) | global, thought disorder | internalizing | 2.2 |
| Cognitive emotion regulation questionnaire (rumination subscale) | internalizing | internalizing | 2.2 |
| Temperament and character inventory (temperaments, harm avoidance subscale) | internalizing | internalizing | 2.3 |

| | | | |
|---|---|---|---|
| NEO five-factor personality inventory (neuroticism factor) | healthy functioning | internalizing | 2.3 |
| Quick inventory of depressive symptomatology (total, sum) | internalizing | internalizing | 2.5 |
| Cognitive failures questionnaire (distractibility subscale) | global, thought disorder | internalizing | 2.5 |
| State-trait anxiety inventory (trait anxiety subscale) | internalizing | internalizing | 2.5 |
| Barratt impulsiveness scale (subscale 2, attentional) | externalizing | internalizing | 2.5 |
| Cognitive emotion regulation questionnaire (self blame subscale) | internalizing | internalizing | 2.5 |
| State-trait anxiety inventory (state anxiety subscale) | internalizing | internalizing | 2.6 |
| Barratt impulsiveness scale (subscale 1, attention) | externalizing | internalizing | 2.7 |
| Ruminative responsiveness scale (total, sum) | internalizing | internalizing | 2.7 |
| Retrospective self-report of inhibition (fear/illness subscale) | internalizing, externalizing | internalizing | 2.7 |
| Perceived stress scale (total) | internalizing | internalizing | 2.7 |
| Montgomery-Asberg depression rating scale (total) | internalizing | internalizing | 2.8 |
| Barratt impulsiveness scale (subscale 1, cognitive instability) | externalizing, thought disorder | internalizing | 2.9 |
| Profile of mood states (anger subscale t-score) | externalizing | internalizing | 2.9 |
| Experiences in close relationships inventory (reassurance subscale) | social, externalizing | internalizing | 3.0 |
| Depression anxiety stress scale (anxiety subscale) | internalizing | internalizing | 3.0 |
| Profile of mood states (tension subscale t-score) | internalizing | internalizing | 3.1 |
| Profile of mood states (depression subscale t-score) | internalizing | internalizing | 3.1 |
| Depression anxiety stress scale (depression subscale) | internalizing | internalizing | 3.2 |
| Profile of mood states (fatigue subscale t-score) | internalizing | internalizing | 3.4 |
| Cognitive emotion regulation questionnaire (catastrophizing subscale) | internalizing | internalizing | 3.4 |
| Experiences in close relationships inventory (pulling back subscale) | social, externalizing | internalizing | 3.4 |
| Clinical global impression (severity score) | global | internalizing | 3.6 |
| Depression anxiety stress scale (stress subscale) | internalizing | internalizing | 3.8 |
| Domain specific risk taking (risk perception, recreational subscale) | externalizing, risk, social, global | externalizing | 1.2 |
| Young mania rating scale (disruptive behavior subscale) | externalizing | externalizing | 1.4 |
| Domain specific risk taking (risk perception, health subscale) | externalizing, risk, social, global | externalizing | 1.5 |
| Domain specific risk taking (risk perception, ethical subscale) | externalizing, risk, social, global | externalizing | 2.3 |
| Temperament and character inventory (characters, cooperativeness subscale) | healthy functioning | externalizing | 2.5 |

| | | | |
|---|---|---|---|
| Young mania rating scale (sleep subscale) | thought disorder, somatic, internalizing | externalizing | 2.7 |
| Temperament and character inventory (temperaments, persistence subscale) | healthy functioning, internalizing | externalizing | 2.9 |
| NEO five-factor personality inventory (agreeableness factor) | healthy functioning | externalizing | 3.0 |
| Multidimensional scale for perceived social support (friend subscale) | social, global, externalizing | externalizing | 3.2 |
| Temperament and character inventory (temperaments, reward dependence subscale) | externalizing, reward | externalizing | 3.5 |
| Young mania rating scale (language, thought disorder subscale) | thought disorder, externalizing | externalizing | 3.5 |
| Panic disorder severity scale (panic frequency subscale) | internalizing | externalizing | 3.7 |
| Temporal experience of pleasure scale (anticipatory subscale) | reward, thought disorder | externalizing | 3.8 |
| Multidimensional scale for perceived social support (family subscale) | social, global, externalizing | externalizing | 4.1 |
| Behavioral inhibition/activation scale (activation, reward responsiveness subscale) | reward, internalizing | externalizing | 4.2 |
| Cognitive emotion regulation questionnaire (refocus/planning subscale) | healthy functioning, externalizing | externalizing | 4.5 |
| Multidimensional scale for perceived social support (significant other subscale) | social, global, externalizing | externalizing | 4.6 |
| Profile of mood states (vigor subscale t-score) | externalizing | externalizing | 4.7 |
| Temperament and character inventory (characters, self directedness subscale) | healthy functioning | externalizing | 4.8 |
| NEO five-factor personality inventory (extraversion factor) | healthy functioning | externalizing | 4.8 |
| NEO five-factor personality inventory (conscientiousness factor) | healthy functioning | externalizing | 4.8 |
| Cognitive emotion regulation questionnaire (positive reappraisal subscale) | healthy functioning | externalizing | 5.6 |
| Young mania rating scale (irritability subscale) | externalizing | externalizing | 6.7 |
| Snaith-Hamilton pleasure scale (total) | reward, externalizing | externalizing | 15.9 |
| Young mania rating scale (speech subscale) | thought disorder, externalizing | cognition | 0.0 |
| Fagerstrom test for nicotine dependence (total) | externalizing | cognition | 0.0 |
| TestMyBrain (fast reaction test) | healthy functioning, cognitive | cognition | 0.1 |
| Domain specific risk taking (risk perception, financial subscale) | externalizing, risk, social, global | cognition | 0.1 |
| TestMyBrain (word sentence association paradigm) | healthy functioning, cognitive | cognition | 0.1 |
| Young mania rating scale (content subscale) | thought disorder | cognition | 0.4 |
| TestMyBrain (reading the mind in the eyes test) | healthy functioning, cognitive | cognition | 0.5 |
| TestMyBrain (matrix reasoning) | healthy functioning, cognitive | cognition | 0.5 |
| Experiences in close relationships inventory (anxiety subscale) | social, externalizing | cognition | 0.5 |
| TestMyBrain (continuous performance test) | healthy functioning, cognitive | cognition | 0.7 |
| Clinical global impression (global improvement score) | global | cognition | 0.7 |
| Young mania rating scale (sexual interest subscale) | thought disorder, externalizing | cognition | 0.9 |

| | | | |
|---|---|---|---|
| Domain specific risk taking (risk perception, social subscale) | externalizing, risk, social, global | cognition | 1.0 |
| TestMyBrain (fast choice test) | healthy functioning, cognitive | cognition | 1.1 |
| Shipley Institute of living scale (vocab subscale t-score) | healthy functioning, cognitive | cognition | 1.4 |
| Shipley Institute of living scale (IQ score) | healthy functioning, cognitive | cognition | 1.4 |
| NEO five-factor personality inventory (openness factor) | healthy functioning | cognition | 1.4 |
| Young mania rating scale (elevated mood subscale) | thought disorder, internalizing | cognition | 1.6 |
| TestMyBrain (symbol matching test) | healthy functioning, cognitive | cognition | 1.6 |
| Young mania rating scale (increased motor subscale) | somatic, externalizing | cognition | 1.7 |
| Young mania rating scale (appearance subscale) | internalizing, thought disorder | cognition | 2.5 |
| Multnomah community ability scale (total) | social, global, externalizing | cognition | 4.5 |
| Domain specific risk taking (risk taking, social subscale) | externalizing, risk, social, global | cognition | 5.2 |
| Temporal experience of pleasure scale (consummatory subscale) | reward, thought disorder | cognition | 5.5 |
| Behavioral inhibition/activation scale (activation, drive subscale) | reward, internalizing | cognition | 6.2 |
| Behavioral inhibition/activation scale (activation, fun seeking subscale) | reward, internalizing | cognition | 6.7 |
| Experiences in close relationships inventory (frustrated subscale) | social, externalizing | cognition | 8.9 |
| Cognitive emotion regulation questionnaire (acceptance subscale) | healthy functioning, internalizing | cognition | 11.0 |
| Cognitive emotion regulation questionnaire (perspective subscale) | healthy functioning | cognition | 14.4 |
| Cognitive emotion regulation questionnaire (positive refocusing subscale) | healthy functioning, thought disorder | cognition | 19.3 |
| Temperament and character inventory (characters, self transcendence subscale) | thought disorder, internalizing, healthy functioning | social/reward | 3.7 |
| Alcohol tobacco caffeine use questionnaire (total) | externalizing | social/reward | 4.6 |
| Domain specific risk taking (risk taking, recreational subscale) | externalizing, risk, social, global | social/reward | 4.7 |
| Barratt impulsiveness scale (subscale 1, motor) | externalizing | social/reward | 5.5 |
| Barratt impulsiveness scale (subscale 1, cognitive complexity) | externalizing, thought disorder | social/reward | 5.8 |
| Barratt impulsiveness scale (subscale 1, perseverance) | externalizing, healthy functioning | social/reward | 6.0 |
| Cognitive emotion regulation questionnaire (blaming others subscale) | externalizing, internalizing | social/reward | 6.6 |
| Temperament and character inventory (temperaments, novelty seeking subscale) | externalizing, reward | social/reward | 7.6 |
| Barratt impulsiveness scale (subscale 2, motor) | externalizing | social/reward | 7.7 |
| Domain specific risk taking (risk taking, health subscale) | externalizing, risk, social, global | social/reward | 9.4 |
| Domain specific risk taking (risk taking, ethical subscale) | externalizing, risk, social, global | social/reward | 12.1 |
| Domain specific risk taking (risk taking, financial subscale) | externalizing, risk, social, global | social/reward | 12.1 |

| | | | |
|---|---|---|---|
| Experiences in close relationships inventory (avoid closeness subscale) | social, externalizing | social/reward | 14.2 |

**Table S1 legend**. These measures (each row) correspond to the x- and y-axes on the individual-differences correlation matrix in main text **Fig. 3A**. A priori labels refer to a pre-mapping of each measure to potential dimensions of functioning based on the literature and consensus across the study team. This was implemented to reduce the possibility of researcher bias in naming clusters (3rd column) that resulted from hierarchical agglomerative clustering of the matrix in **Fig. 3A**. Once the clustering solution was optimized (see **Methods**), PCA was performed on each cluster of measures, and based on the relative importance of each measure's factor loading onto the first PC (4th column), we used a priori labels to guide naming conventions for each cluster. Avg. = average. Note that measures within each cluster are sorted in ascending order of factor loading. See [1] for more detailed information about session acquisition, source references, versioning, and quality assurance analyses for the TCP dataset.

### Supplementary Figures

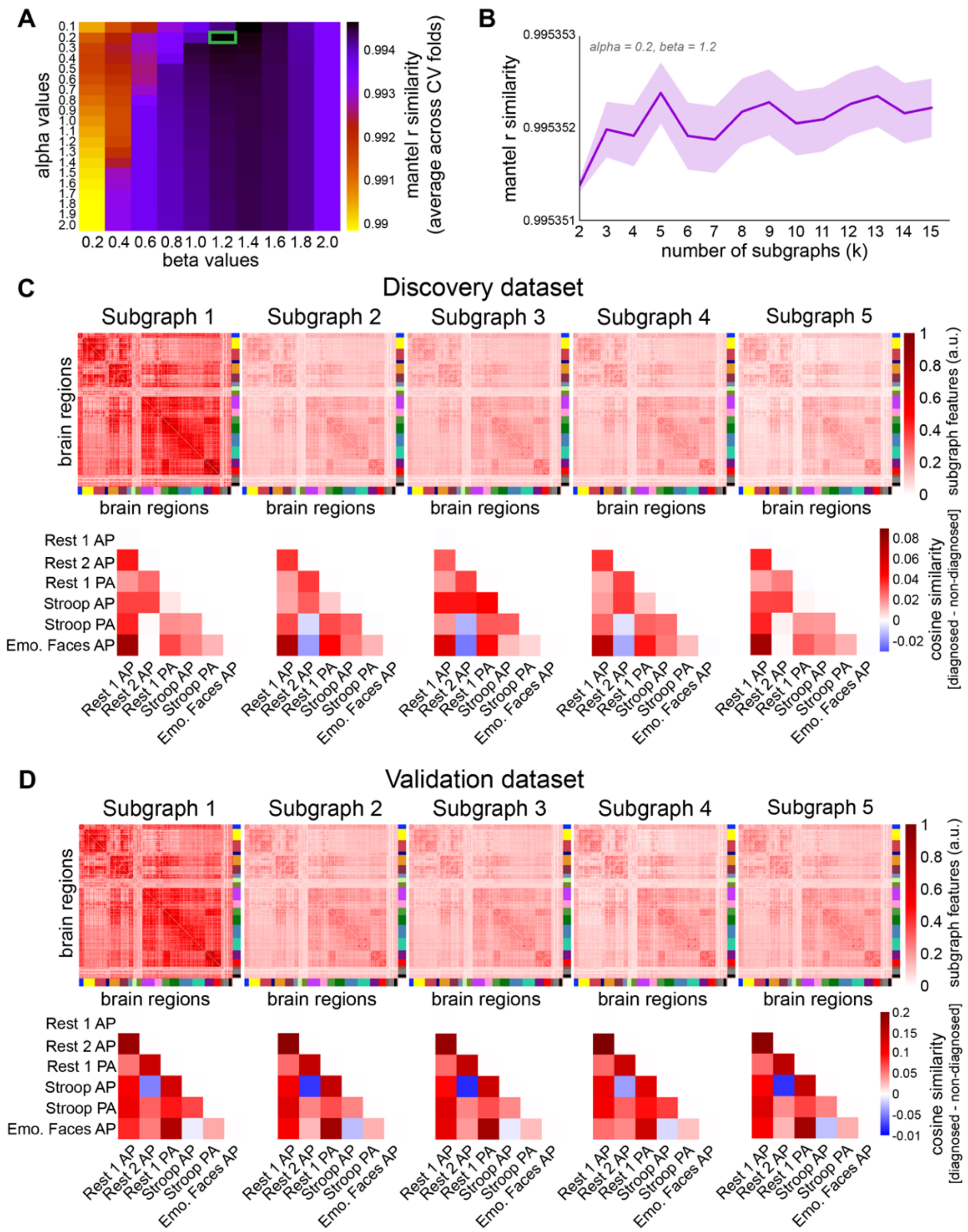


**Figure S1. Validation and optimization of non-negative matrix factorization (NMF) for uncovering brain network dynamics.** (**A**) Cross validation results for tuning NMF parameters alpha and beta. Mantel *r* estimates a nonparametric correlation between the original FC input and the NMF-reconstructed FC. The optimal parameter pair is highlighted in green. (**B**) Same as A, but for the number of subgraphs (*k*), while holding alpha and beta at the optimal levels in A. Cross-validation accuracy was stable for 5-10 subgraphs, and we chose a solution of k=5 for consistency with prior literature[2-5].(**C**) *Top*: NMF subgraph features, reformatted into region-by-region networks. *Bottom*: main-text **Fig. 2F**, including all subgraphs. (**D**) Same as C (discovery dataset), except for the validation dataset.

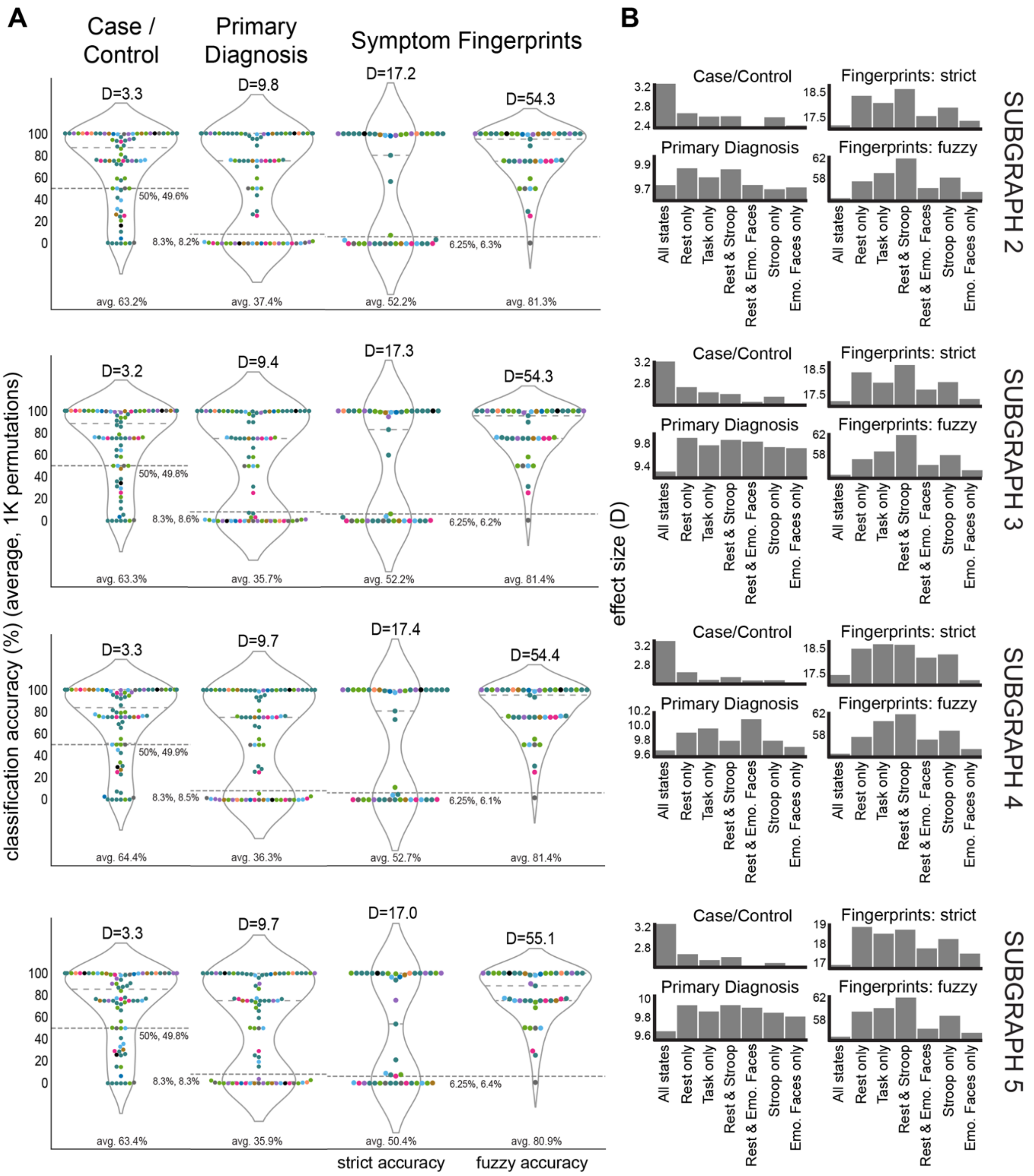


**Figure S2. Brain network dynamics can accurately separate symptom fingerprints, subgraphs 2 through 5 (paneled rows).** (**A**) Classification accuracies closely match main text **Fig. 4**. Effect sizes of each model (*D*) are listed on the top, along with theoretical chance and empirical chance near dashed grey lines. Average model accuracy across 1000 permutations is listed on the bottom of each plot. (**B**) Following main text **Fig. 5**, case-control classification accuracy varies with amount of connectivity data inputted to NMF; primary diagnoses are classified with varying patterns across models; and fingerprints are best separated by brain network dynamics across states involving shifts from resting-state to Stroop inhibitory cognitive control task.

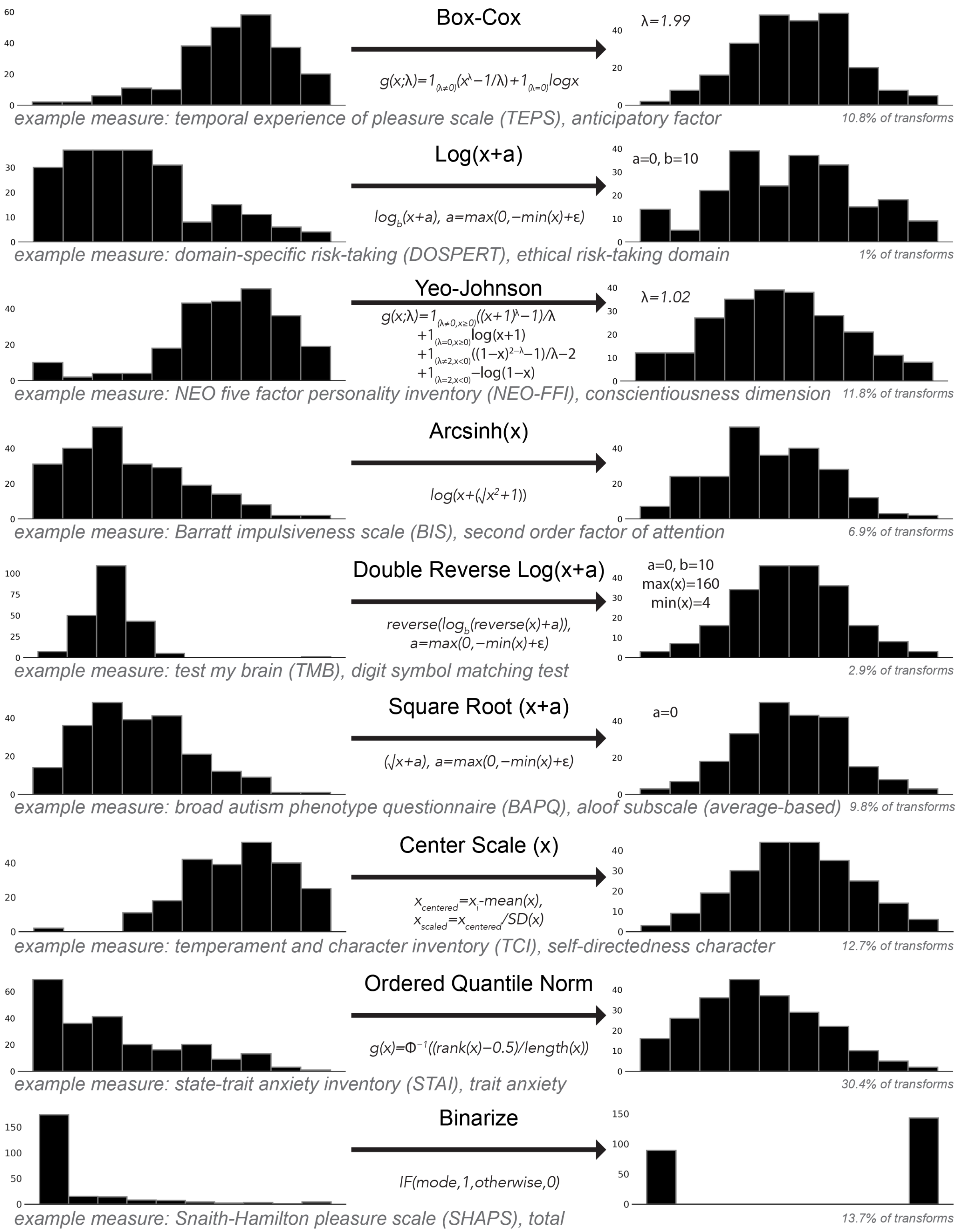


**Figure S3. Normalizing transformations applied to behavioral, cognitive, and clinically-relevant measures for the symptom fingerprinting pipeline.** Using the R package bestNormalize[6], the shown transformation-estimating functions were used on select behavioral measures. In each case, an example implementation is shown along with mathematical formula to demonstrate the transformation process.

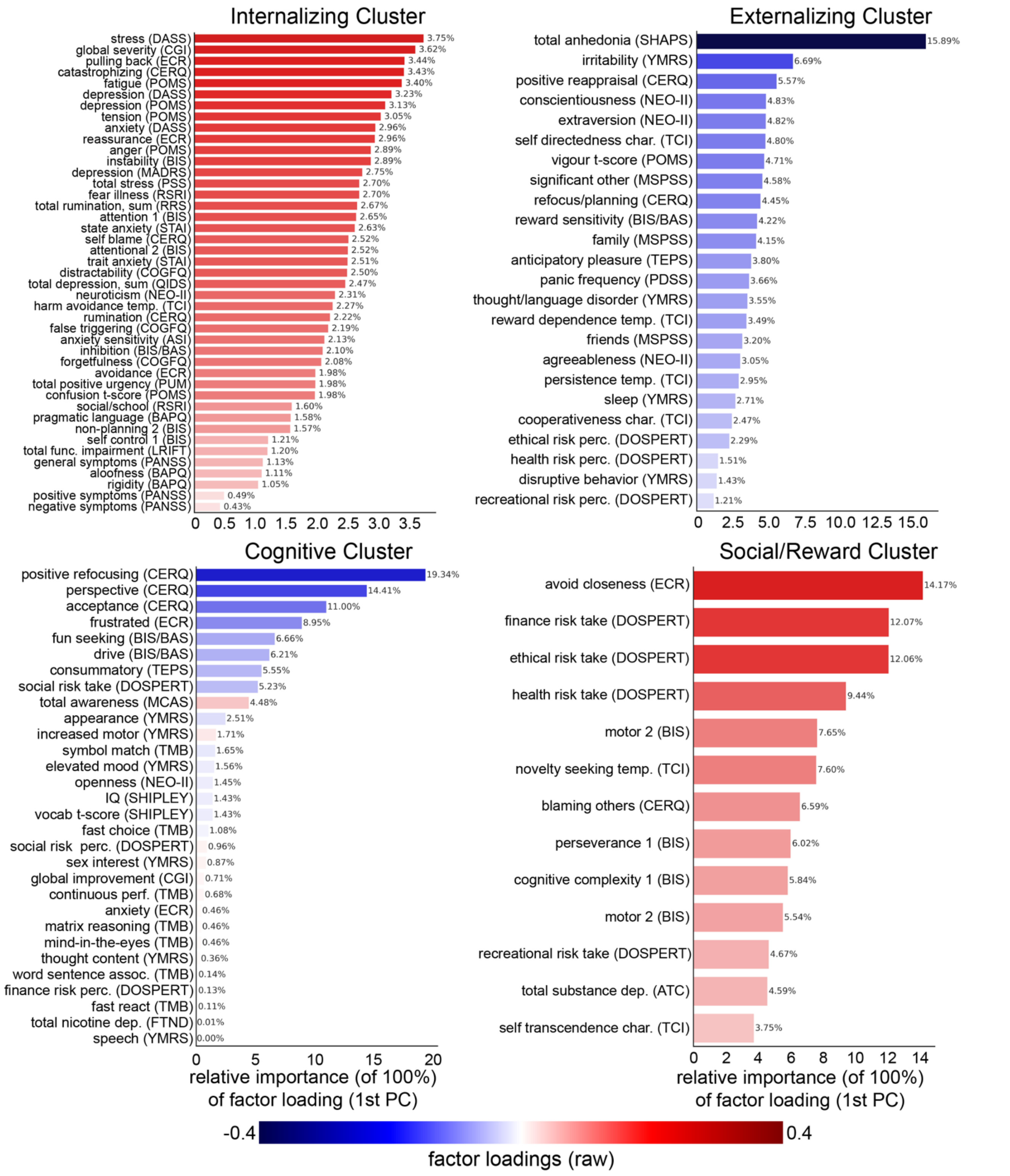


**Figure S4. Uncovering dimensional symptom profiles: expanded PCA loadings.** Corresponding to Results **Fig. 3**, all of the measures in each cluster are shown as well as the relative importance of their factor loadings in the first PC of each cluster. Scale and subscale names (y-axes) are expanded in **Supplemental Table S1,** along with *a priori* domain mapping technique.

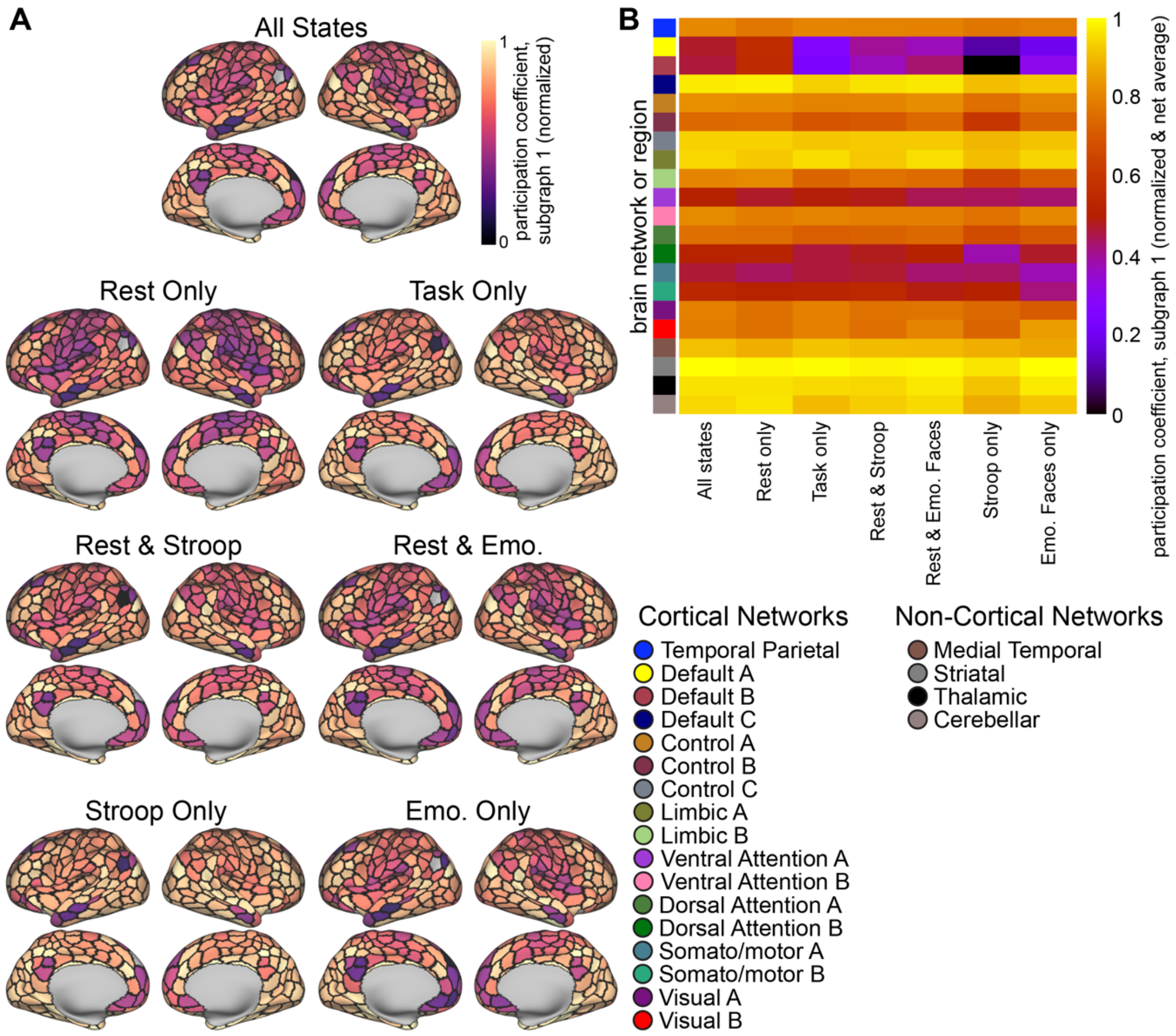


**Figure S5. Hub structure underlying connectivity shifts across varied cognitive states.** To explore the extent that dynamic network reconfigurations are linked with flexible processing, we applied a graph metric called participation coefficient[7], which quantifies hubs, or, how diversely connected a region is to other regions, and has been linked with integrative processes[8,9]. Applying participation coefficient to the first NMF-uncovered subgraph quantifies the extent that features underlying across-state network dynamics exhibit hub properties. (**A**) Participation coefficient (PC) of the first NMF-uncovered subgraph underlying across-state brain network dynamics, with FC estimates from different combinations of states used as inputs (i.e., the underlying network structure for the inputs in **Fig. 5** model comparisons). PC estimates were min-max normalized to a common scale of 0-1. While gross patterns were similar, regions in control network A had increased PC. (**B**) Same as panel B, but functional network averages shown. Select networks exhibited hub properties consistently across all models, such as striatal areas and control (frontoparietal) A networks, and others with more variability, such as default networks A and B. Altogether, network features constraining information processing dynamics were exhibited with different degrees of variability or stability across ranging shifts in cognitive context, depending on the functional brain system.

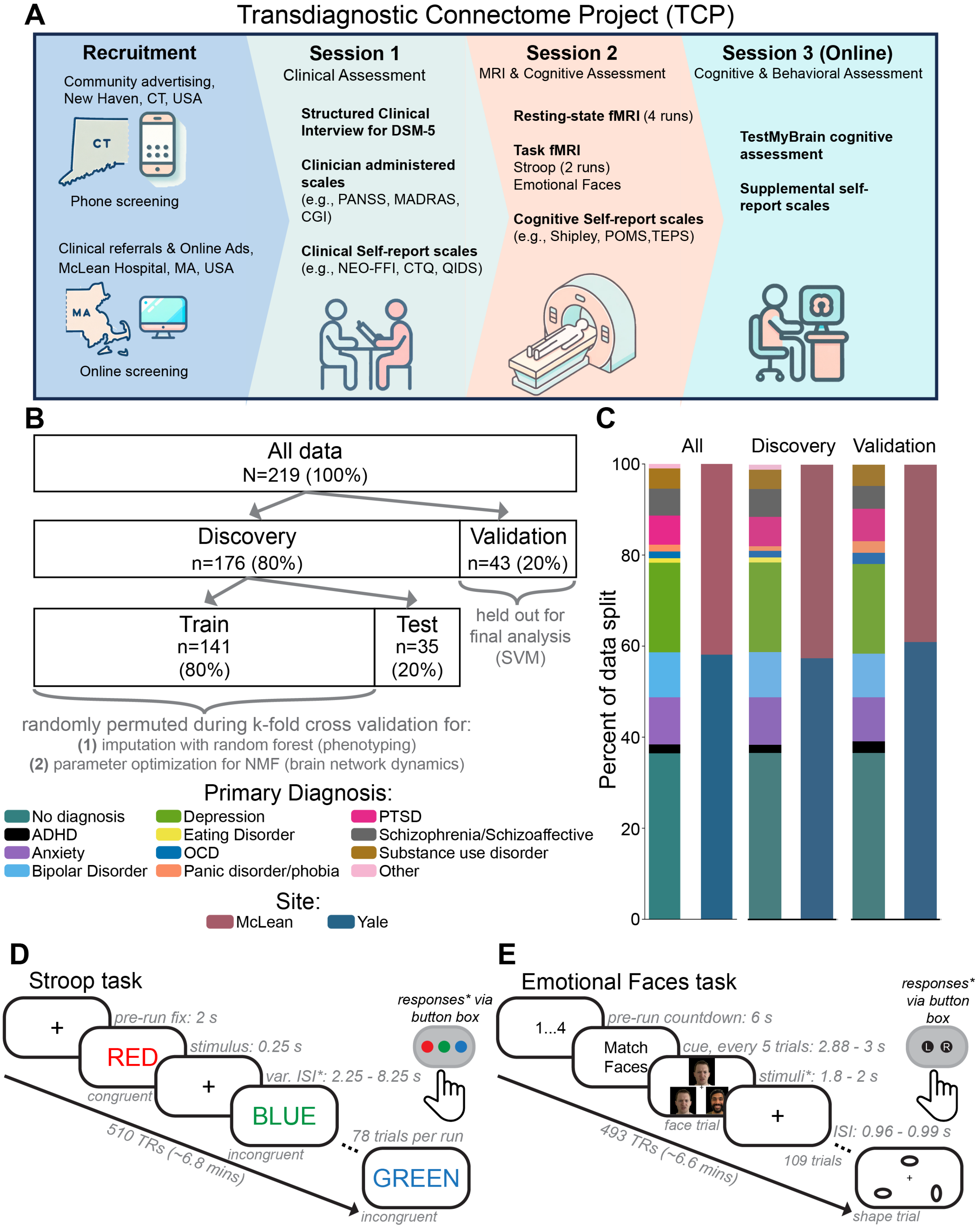
A
Transdiagnostic Connectome Project (TCP)
Recruitment
Community advertising, New Haven, CT, USA
CT
Phone screening
Clinical referrals & Online Ads, McLean Hospital, MA, USA
MA
Online screening
Session 1
Clinical Assessment
Structured Clinical Interview for DSM-5
Clinician administered scales (e.g., PANSS, MADRAS, CGI)
Clinical Self-report scales (e.g., NEO-FFI, CTQ, QIDS)
Session 2
MRI & Cognitive Assessment
Resting-state fMRI (4 runs)
Task fMRI
Stroop (2 runs)
Emotional Faces
Cognitive Self-report scales (e.g., Shipley, POMS,TEPS)
Session 3 (Online)
Cognitive & Behavioral Assessment
TestMyBrain cognitive assessment
Supplemental self-report scales
B
All data
N=219 (100%)
Discovery
n=176 (80%)
Validation
n=43 (20%)
held out for final analysis (SVM)
Train
n=141
(80%)
Test
n=35
(20%)
randomly permuted during k-fold cross validation for:
(1) imputation with random forest (phenotyping)
(2) parameter optimization for NMF (brain network dynamics)
Primary Diagnosis:
No diagnosis
ADHD
Anxiety
Bipolar Disorder
Depression
Eating Disorder
OCD
Panic disorder/phobia
PTSD
Schizophrenia/Schizoaffective
Substance use disorder
Other
Site:
McLean
Yale
C
All
Discovery
Validation
Percent of data split
100
80
60
40
20
0
D
Stroop task
+
pre-run fix: 2 s
RED
stimulus: 0.25 s
congruent
+
var. ISI*: 2.25 - 8.25 s
BLUE
incongruent
78 trials per run
GREEN
incongruent
510 TRs (~6.8 mins)
responses* via button box
E
Emotional Faces task
1...4
pre-run countdown: 6 s
Match Faces
cue, every 5 trials: 2.88 - 3 s
stimuli*: 1.8 - 2 s
face trial
+
ISI: 0.96 - 0.99 s
109 trials
shape trial
493 TRs (~6.6 mins)
responses* via button box
L
R

93

**Figure S6. The Transdiagnostic Connectome Project (TCP) dataset**. (**A**) Example behavioral measures, types of fMRI data, and locale information is listed for each of the three TCP sessions. (**B**) Data were semi-randomly split (constraints to preserve proportionality of primary diagnosis and collection site) into discovery (n=176) and validation groups (n=43). Discovery data was randomly split into training and test sets for various supervised learning and/or parameter optimization steps of the pipeline, and validation data was not used until the final analyses. This ensured that the held-out testing set in classification (validation) did not "leak" information into the training set (discovery). (**C**) Percentages of participants in each primary diagnosis (as given by the SCID-V-RV) and from each collection site in the total, discovery, and validation datasets. Select primary diagnoses were collapsed based on common symptomatology, for example: substance use disorder (SUD) included cocaine use disorder and alcohol use disorder. ADHD = attention-deficit/hyperactivity disorder; OCD = obsessive compulsive disorder; PTSD = posttraumatic stress disorder. (**D**) The Stroop task paradigm used in task fMRI. (**E**) The Emotional Faces task paradigm used in task fMRI Var. ISI = variable interstimulus interval; TR = repetition time (1 TR = 0.8 s in TCP fMRI data). Panels A, D, and E were reproduced with permission from [[1]].
